# Glioma-associated tertiary lymphoid structures are sites of lymphocyte clonal expansion and plasma cell formation

**DOI:** 10.1101/2024.07.04.602038

**Authors:** Pinar Cakmak, Jennifer H. Lun, Aakanksha Singh, Jadranka Macas, Jonathan Schupp, Miriam Köhler, Tatjana Starzetz, Michael C. Burger, Eike Steidl, Lucie Marie Hasse, Elke Hattingen, Karl H. Plate, Yvonne Reiss, Katharina Imkeller

## Abstract

Adult-type diffuse gliomas, the most common primary brain tumors, pose significant clinical challenges due to limited treatment options, restricted anti-tumor immune response and dismal patient prognosis. In this study, we elucidate the immunological function and clinical relevance of intra-tumoral tertiary lymphoid structures (TLS) in adaptive anti-glioma immunity. We conducted a comprehensive, unbiased analysis of lymphoid aggregation in 642 gliomas using a multi-modal approach that combines RNA sequencing with spatial transcriptome and proteome profiling. Our findings reveal that TLS are present in 15% of tumors and correlate with improved overall survival. Gliomas with TLS exhibit a remodeled perivascular space, marked by transcriptional upregulation and spatial redistribution of collagens associated with barrier functions. Furthermore, we demonstrate that TLS maturation into sites of dynamic adaptive immune responses, characterized by clonal T and B cell expansion and IgA+ and IgG+ plasma-cell formation, is driven by efficient early T cell recruitment to the perivascular space.

## Introduction

The advent of immune checkpoint therapy has revolutionized cancer treatment in several solid cancers^1^. Tumors residing within the central nervous system (CNS) however remain a clinical challenge and often represent a difficult target for immunotherapy^2^. Hallmarks of CNS neoplasms are brain-resident cell types and a restrictive blood-brain barrier (BBB) with distinctive extracellular matrix (ECM) components that interfere with immune cell infiltration and drug delivery into the tumor parenchyma^2,3^. Among primary CNS tumors, adult-type diffuse gliomas classified based on the presence or absence of isocitrate dehydrogenase (IDH) mutations, are most common^4,5^. Due to its aggressive, highly invasive and angiogenic growth pattern, the most frequent malignant adult-type diffuse glioma, IDH wildtype glioblastoma (GBM), is currently incurable with a median patient survival of less than 21 months^5–7^. The standard treatment for IDH wildtype GBM includes gross surgical resection, followed by radiotherapy and adjuvant chemotherapy^8^. Attempts to incorporate immune checkpoint inhibition in glioma therapy have largely failed^5,9^, partly due to T cell exclusion and immunosuppressive mechanisms^10–13^.

Intratumoral immune aggregates, such as tertiary lymphoid structures (TLS), have recently gained interest in the field of cancer immunotherapy as they seem to mediate sustained anti-tumor immune responses^14–21^. TLS are ectopically formed aggregates of lymphoid and stromal cells, comprising T cell zones with antigen-presenting dendritic cells, and B cell zones with germinal centers^19–21^. TLS support local immune responses through several mechanisms, including antigen presentation to T cells^22,23^ or differentiation of B cells into plasma cells that secrete tumor-specific antibodies^24,25^. These roles in adaptive immunity make TLS a promising biomarker for patient stratification in immune checkpoint therapy of various cancer types including non-small-cell lung cancer^26,27^.

TLS have recently also been identified in adult-type diffuse gliomas^28,29^. However, the functionality of these structures in anti-glioma immunity and their association with patient survival remain unknown. Given the immune privilege of the brain and the immunosuppressive nature of the glioma tumor microenvironment (TME), we aimed to systematically study the prevalence and clinical relevance of TLS in a large glioma patient cohort (N=642) and determine the spatial architecture and immunological function of glioma-associated TLS.

Our study presents the first evidence for germinal center-like reactions and plasma cell formation within glioma-associated TLS, indicative for their immunological function, which is associated with improved patient survival. Through multi-modal spatial profiling, we reveal the non-canonical expression of ECM components and remodeling of the glioma vasculature that precedes TLS formation and maturation. Moreover, we demonstrate that the maturation of TLS into sites of dynamic adaptive immune responses is shaped by the TME and facilitated by early T cell recruitment into the perivascular space.

## Results

### Prevalence and clinical relevance of TLS in adult-type diffuse glioma

The prevalence and clinical relevance of glioma-associated TLS was determined by screening a cohort of 452 IDH-wildtype (wt) and 190 IDH-mutant (mut) adult-type diffuse gliomas, using an IHC-based imaging approach (**Fig. 1A and Fig. S1A**). FFPE-tissue sections (n=642) were screened for CD20+ B cells. Lymphoid aggregates with a minimum of 50 cells, including CD20+ B cells, were considered TLS and further analyzed (**Fig. 1A**). Our screening revealed 15% of gliomas with at least one TLS, which we classified as TLS+, whereas all other tumors were classified as TLS-. TLS were localized to both, the tumor parenchyma and the leptomeninges, and were more commonly found in proximity to leptomeninges (**Fig. S1B and S1C**). A higher rate of meningeal enhancement was observed in tumors with at least one leptomeningeal TLS, suggesting a significant historadiological correlation between meningeal enhancement and TLS (**Fig. S1D**).

**Figure 1:**
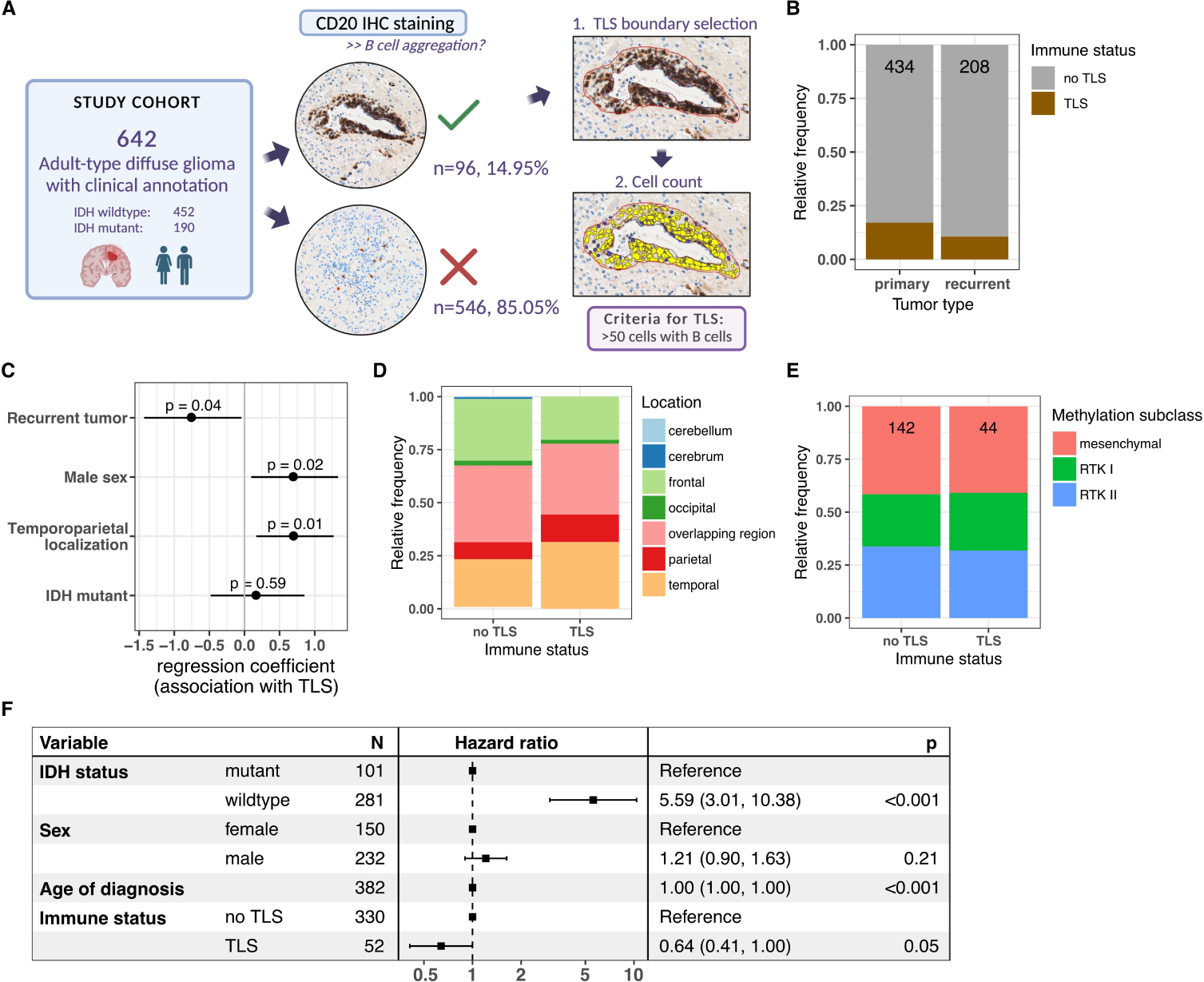
The presence of TLS in adult-type diffuse glioma is associated with improved overall survival. **A:** Schematic representation of the screening approach for TLS identification in adult-type diffuse glioma. Tumor specimens (N=642) were subjected to anti-CD20 immuno-histochemistry (IHC) staining. Tertiary lymphoid structures (TLS) were defined as lymphoid aggregations of a minimum of 50 cells with the presence of CD20+ B cells. **B:** Relative frequency of TLS+ tumors in primary (n=434) and recurrent (n=208) adult-type diffuse gliomas. **C:** Results of logistic regression to determine the association of different tumor characteristics (primary/recurrent, patient sex, tumor location, IDH mutational status) with the presence of TLS. Dots represent the regression coefficients, and lines represent standard errors. **D:** Relative frequency of tumor location in patients with TLS+ and TLS-adult-type diffuse gliomas. Colors indicate tumor location within the brain. **E:** Relative frequency of methylation subclasses in IDH-wt glioblastoma patients with TLS-(N=142) and TLS+ (N=44) tumors. Colors indicate methylation subclasses. **F:** Forest plot depicting the hazard ratio (HR) and 95% CI of the association of overall survival with different clinical parameters (tumor IDH mutational status, patient sex, patient age at diagnosis, TLS+/TLS-status). A multinomial Cox proportional hazard regression model was applied to determine statistical significance.

Our specimen collection included 434 primary and 208 recurrent gliomas with a slightly higher prevalence of TLS in primary tumors compared to recurrent tumors (**Fig. 1B**). By applying multinomial logistic regression, we determined that the presence of TLS was independent of the IDH mutation status (p=0.57), negatively associated with tumor recurrence (p=0.03), and positively associated with male biological sex (p=0.02) and temporoparietal tumor location (p=0.01) (**Fig. 1B-D, S1E and S1F**). For IDH-wt tumors, the prevalence of TLS was independent of the methylation-based classification into mesenchymal, RTKI and RTKII glioblastoma subtypes (**Fig. 1E**). Survival data and follow-up data were available for 383 patients. Using multinomial Cox proportional hazard regression analysis, we identified TLS status as a positive predictor for overall survival and progression-free survival independent of IDH mutational status, biological sex and age at diagnosis (**Fig. 1F and Fig. S1G**).

TLS formation in other tumor entities has been associated with high tumor mutational burden (TMB) leading to the availability of tumor-specific antigens^30^. However, when comparing the TMB values between TLS+ and TLS-gliomas, no significant difference was observed (**Fig. S1H**). Three tumors (2 TLS+, 1 TLS-) showed hypermutation with a TMB value ≥ 17 mut/Mb.

### TLS+ gliomas are enriched for transcripts reminiscent of vascular remodeling and depleted for neuronal signatures

We next performed RNA sequencing, followed by differential gene expression and gene set enrichment analysis to compare the transcriptional profiles of TLS+ (n=44) and TLS-(n=20) adult-type diffuse gliomas (**Fig. 2 and Fig. S2**). TLS-tumors were characterized by higher expression of transcripts related to neuronal identity and synaptic function, independent of the IDH mutational status of the tumor or patients’ biological sex (**Fig. 2A-2C**). Transcripts up-regulated in TLS-tumors included *SYT4, TNR* and *RAPGEF* (**Fig. 2A and 2B, Supplementary table 1**), which are expressed at high levels by neurons according to a single-cell reference map of the glioblastoma TME^31^ (**Fig. 2D**). Indeed, cell type deconvolution with this brain tumor specific reference revealed an overall enrichment of cells from neural lineages (neurons, radial glial cells, oligodendrocyte precursor cells, oligodendrocytes, astrocytes) in the TLS-tumors (**Fig. 2E**).

**Figure 2:**
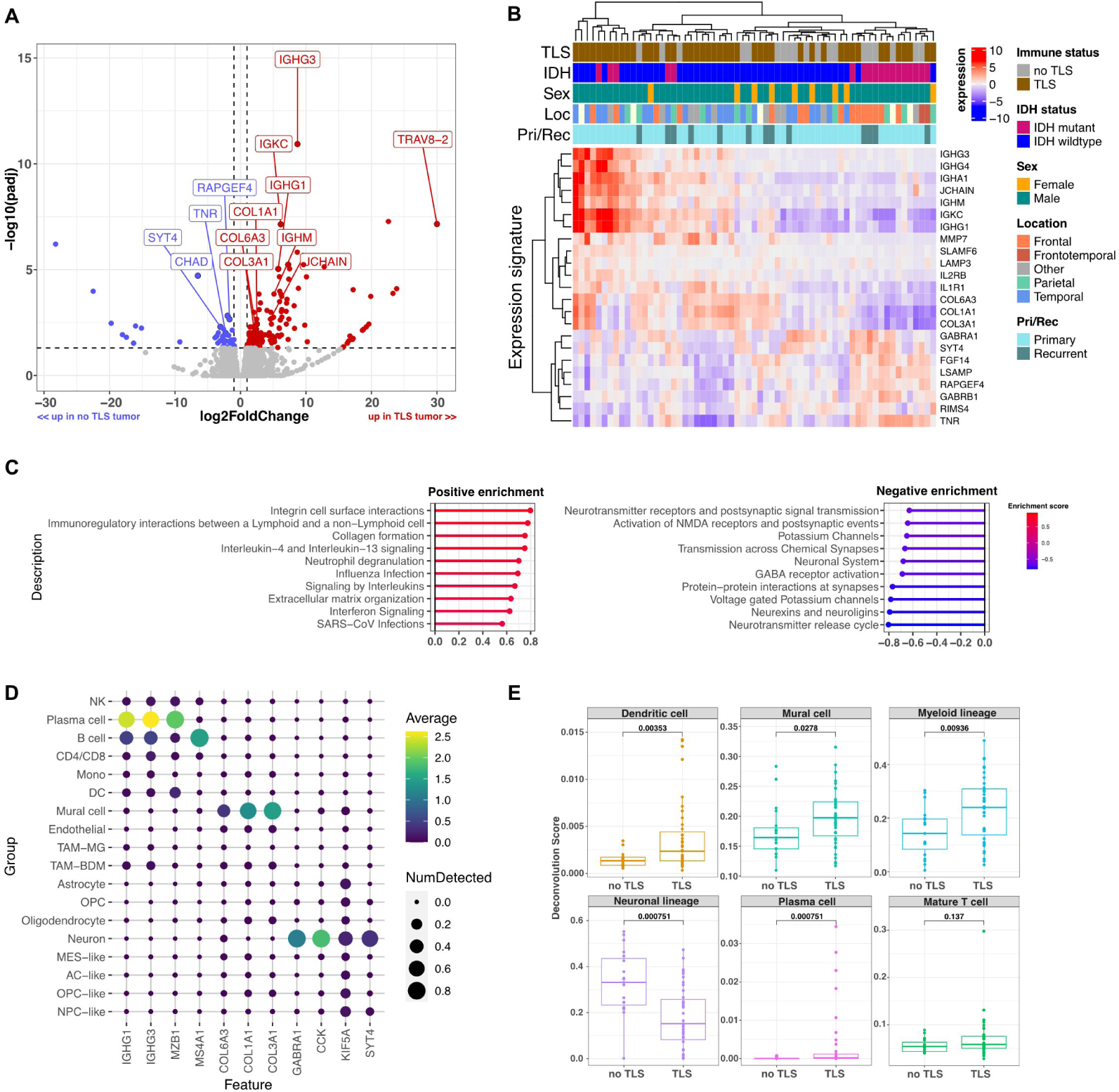
TLS+ gliomas are enriched for transcripts reminiscent of vascular remodeling and depleted for neuronal signatures. **A:** Volcano plot showing differentially expressed genes between TLS+ and TLS-tumors. Labeled genes had an absolute logarithmic fold change > 2 and adjusted p-value < 0.05. Gene expression measured by bulk tumor RNA sequencing of TLS+ (n=44) and TLS-(n=20) gliomas. See also Fig. S2 for differential gene expression analysis. **B:** Heatmap displaying the expression of selected up- and down-regulated genes across all sequenced samples. Colored column annotations indicate IDH-mutational status, patient biological sex, immune status, location and primary/recurrent status of tumors. Expression values are normalized and scaled. **C:** Gene set enrichment analysis (GSEA) results based on differential gene expression between TLS+ and TLS-tumors. Pathways with positive enrichment score (higher expressed in TLS+) are shown in red, negatively enriched pathway in blue. **D:** Expression of specific marker genes in different cell types of a glioblastoma single-cell reference atlas (GBMmap^31^). The color indicates the average normalized expression of a gene (column) in each of the cell type (rows). The dot size indicates the fraction of cells in which the gene was detected. **E:** Transcriptional deconvolution score of different cell types between TLS- and TLS+ tumors. SpatialDecon^56^ with a glioblastoma specific single-cell reference was used for transcriptional deconvolution^31^. The deconvolution score represents the relative frequency of a cell type, ranging from 0 to 1. P-values are determined by the Wilcoxon test.

In contrast, TLS+ tumors showed higher levels of transcripts typically expressed by B cells and plasma cells, including immunoglobulin gene segments such as *IGHG3* and *IGKV1-5*, as well as transcripts linked to interferon and interleukin signaling (*IL2RB* and *IL1R1*) (**Fig. 2A-2D, Supplementary table 1**). Transcriptional cell type deconvolution revealed an overall enrichment of immune cells, including dendritic cells, plasma cells and myeloid cells, in the TLS+ tumors (**Fig. 2E**).

In addition, TLS+ tumors expressed higher levels of transcripts related to collagen matrix, such as *COL6A1*, *COL1A1* and *COL3A1*, and extracellular matrix remodeling, e.g. *MMP7* (**Fig. 2A-2C, Supplementary table 1**). Transcriptional cell type deconvolution indicated an enrichment of mural cells in TLS+ tumors (**Fig. 2E**), which also constitute the main source of collagen expression in gliomas according to the single-cell reference atlas^31^ (**Fig. 2D**). Of note, a fraction of these collagens (e.g. Col6) are expressed by specific fibroblast subtypes associated with the brain vasculature in high grade glioma, which are absent in normal brain^32,33^. The remodeling of extracellular matrix in TLS+ tumors was confirmed by highplex immunofluorescence imaging, where we observed restructuring of the Col4+ basal lamina around vessels, which constitutes an element of the blood-brain barrier (**Fig. S3A-S3C**). Interestingly, we identified Col4 fragments within CD163+ macrophages, suggestive of an uptake of extracellular matrix components by tumor-associated macrophages^34^ (**Fig. S3B and S3C**). The immunofluorescence stainings also illustrated the expression of Col6 localized around intra-tumoral blood vessels (**Fig. S3A-S3C**).

### Transcriptional signatures of TLS indicate differences in cellular composition

In order to characterize the cellular and molecular heterogeneity of glioma-associated TLS, we applied a multi-modal spatial profiling approach consisting of spatial transcriptome profiling (whole transcriptome, GeoMx DSP), whole slide multiplex immunofluorescence staining (6 markers, LabSat and VectraPolaris) and highplex immunofluorescence staining (32 markers, COMET) to consecutive sections of the same tissue specimen (**Fig. 3A**). This approach allowed us to trace single TLS through the different molecular layers.

**Figure 3:**
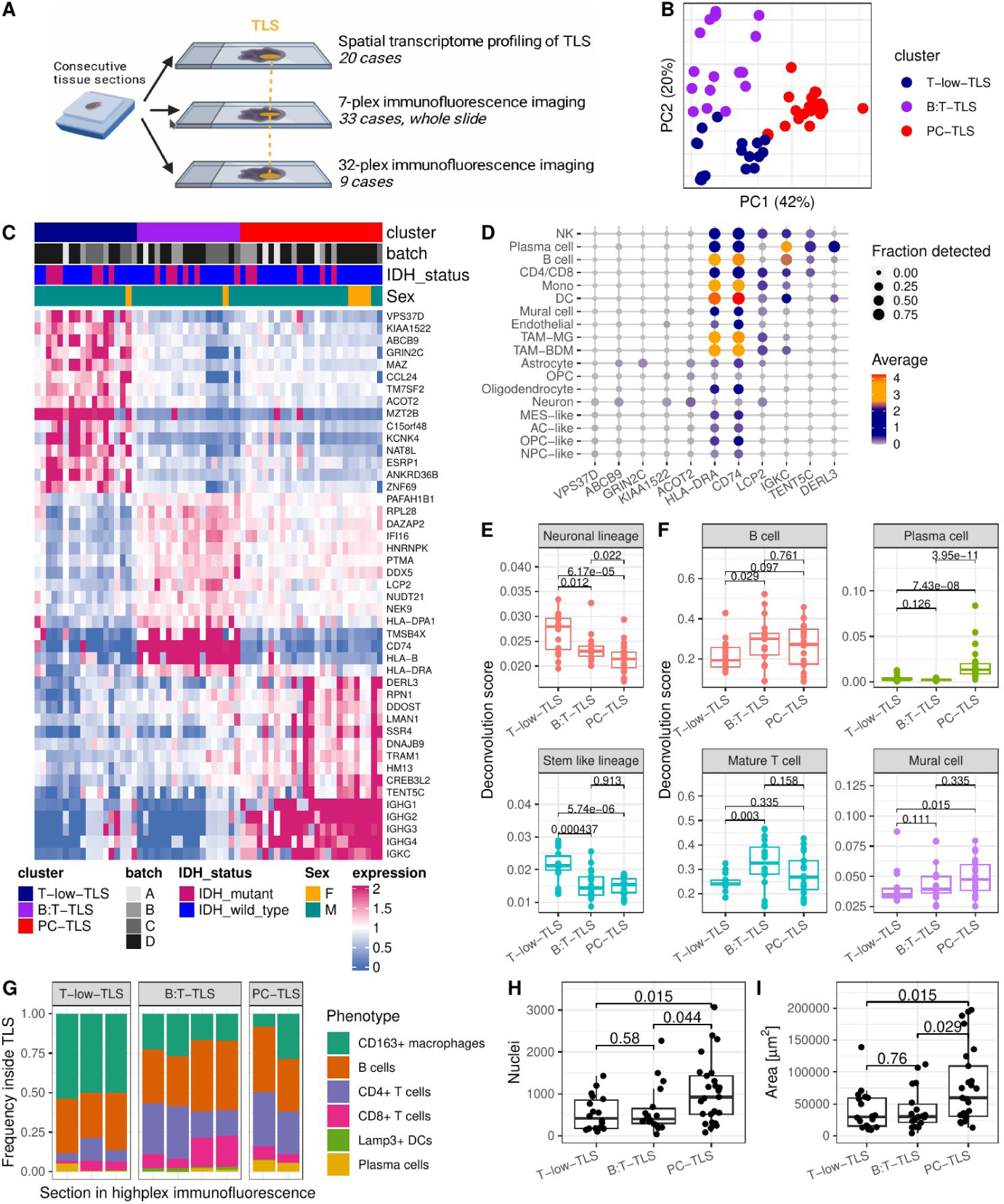
Transcriptional signatures of TLS indicate differences in cellular composition. **A**: Schematic representation of the multi-omic spatial profiling approach based on consecutive tissue sectioning. **B**: PCA analysis of transcriptional profiles from TLS regions (spatial transcriptome profiling with GeoMx). TLS segregate/cluster in three different subtypes with distinct transcriptional profiles (T-low-TLS, B:T-TLS, PC-TLS) are highlighted in blue, purple and red respectively. The percentages of variance explained by each PC are indicated. See also Fig. S4 for PCA results. **C**: Heatmap showing normalized expression of the 15 most important marker genes for the three different TLS subtypes. Each column represents one TLS, the columns are arranged according to TLS subtypes (top annotation; T-low-TLS, B:T-TLS, PC-TLS). Experiment batch (i.e. experiment run), IDH mutational status and patient biological sex are indicated in the top annotations. **D**: Expression of specific marker genes in different cell types of a glioblastoma single-cell reference atlas (GBM map). The color indicates the average normalized expression of a gene (column) in each of the cell types (rows). The dot size indicates the fraction of cells in which the gene was detected. **E and F:** Transcriptional deconvolution score of different tumors (E) and immune /stromal (F) cell types between the three TLS subtypes. SpatialDecon with a glioblastoma specific single-cell reference was used for transcriptional deconvolution. The deconvolution score represents the relative frequency of a cell type, ranging from 0 to 1. P-values are determined by the Wilcoxon test. **G**: Relative frequencies of immune cell phenotypes detected by highplex imaging within the three TLS subtypes. Colors correspond to macrophages (CD163_total), B cells (CD20_total), CD4+ T cells (CD4_total), CD8+ T cells (CD8_total), dendritic cells (Lamp3_total) and plasma cells (PC_total). For the marker definition used to define cell types see Supplementary Table 2. **H and I**: Nuclei counts (H) and areas (I) within the CD20+ TLS regions of interest profiled by spatial transcriptomics. TLS are grouped according to TLS subtype (x-axis). P-values in H and I were calculated using Wilcoxon test.

The transcriptional profiles of 61 different TLS from 20 different tumor samples were investigated using spatial transcriptome profiling. Between one and six TLS per sample were selected based on CD20+ immunofluorescence staining, and an adjacent CD20-negative tumor rich region was included as control. Immune cell transcripts such as *MS4A1*, *CD19*, *CD3D* and *PTPRC* were expressed 2-3 times higher in TLS areas compared to the tumor control regions (**Fig. S4A**).

We applied dimension reduction and clustering to the transcriptional profiles of TLS areas and thereby delineated three subtypes of TLS with distinct transcriptional profiles (**Fig. 3B and 3C; Fig. S4B and S4C**). The first TLS subtype, which we named *T-low-TLS* (low T cell abundance), was characterized by the expression of transcripts preferentially expressed by neurons or astrocytes according to the glioblastoma single-cell reference atlas, such as *ACOT2*, *GRIN2C* and *ABCB9* (**Fig. 3C and 3D**). Transcriptional cell type deconvolution also indicated that tumor cells of the neuronal-like and stem-like lineages were enriched in T-low-TLS (**Fig. 3E**).

The second TLS subtype, named *B:T-TLS* (interactions of B and T cells), was characterized by the expression of genes related to antigen presentation (*CD74*, *HLA-B*, *HLA-DRA*), which are predominantly expressed by macrophages, B cells and dendritic cells (**Fig. 3C and 3D**). The third TLS subtype, denominated *PC-TLS* (plasma cell rich), showed higher levels of genes expressed by plasma cells, e.g. immunoglobulin gene segments, *DERL3* or *TENT5C* (**Fig. 3C and 3D**). Transcriptional cell type deconvolution indicated that B cells, T cells and mural cells were more abundant in B:T-TLS and PC-TLS compared to T-low-TLS (**Fig. 3F**). When matching the TLS subtype annotation from spatial transcriptomics to the highplex immunofluorescence data, B:T-TLS and PC-TLS showed higher proportions of CD4+ and CD8+ T cells compared to T-low-TLS, whereas CD163+ macrophages were enriched in T-low-TLS (**Fig. 3G**). PC-TLS had higher nuclei counts and occupied larger areas compared to T-low-TLS and B:T-TLS (**Fig. 3H and 3I**). No significant association was found between leptomeningeal TLS location and TLS subtype (**Fig. S4E)** or between or IDH mutation status and TLS subtype (**Fig. 3C**).

### B:T-TLS and PC-TLS show evidence for dynamic adaptive immune responses

TLS have been described to form separated T and B cell areas (zonation), similar to functional regions described in lymph nodes^35^. Since the TLS in our glioma samples showed considerable amounts of T cells and plasma cell formation, we next investigated whether they also showed signs of germinal center like reactions (**Fig. 4**). We applied multiplex immunofluorescence staining and revealed that the average distance between T cells and their closest B cell was slightly higher in B:T-TLS and PC-TLS when compared to T-low-TLS (**Fig. 4A and 4B**). The percentage of T cells within 15 µm distance of B cells was slightly lower in B:T-TLS and PC-TLS compared to T-low-TLS (**Fig. 4C**). These findings suggest, that T cells and B cells are more intermixed in T-low-TLS, whereas they tend to accumulate in separate regions in B:T-TLS and PC-TLS, as illustrated in a representative density heatmap of B and T cells within the three different subtypes of TLS (**Fig. 4D**).

**Figure 4:**
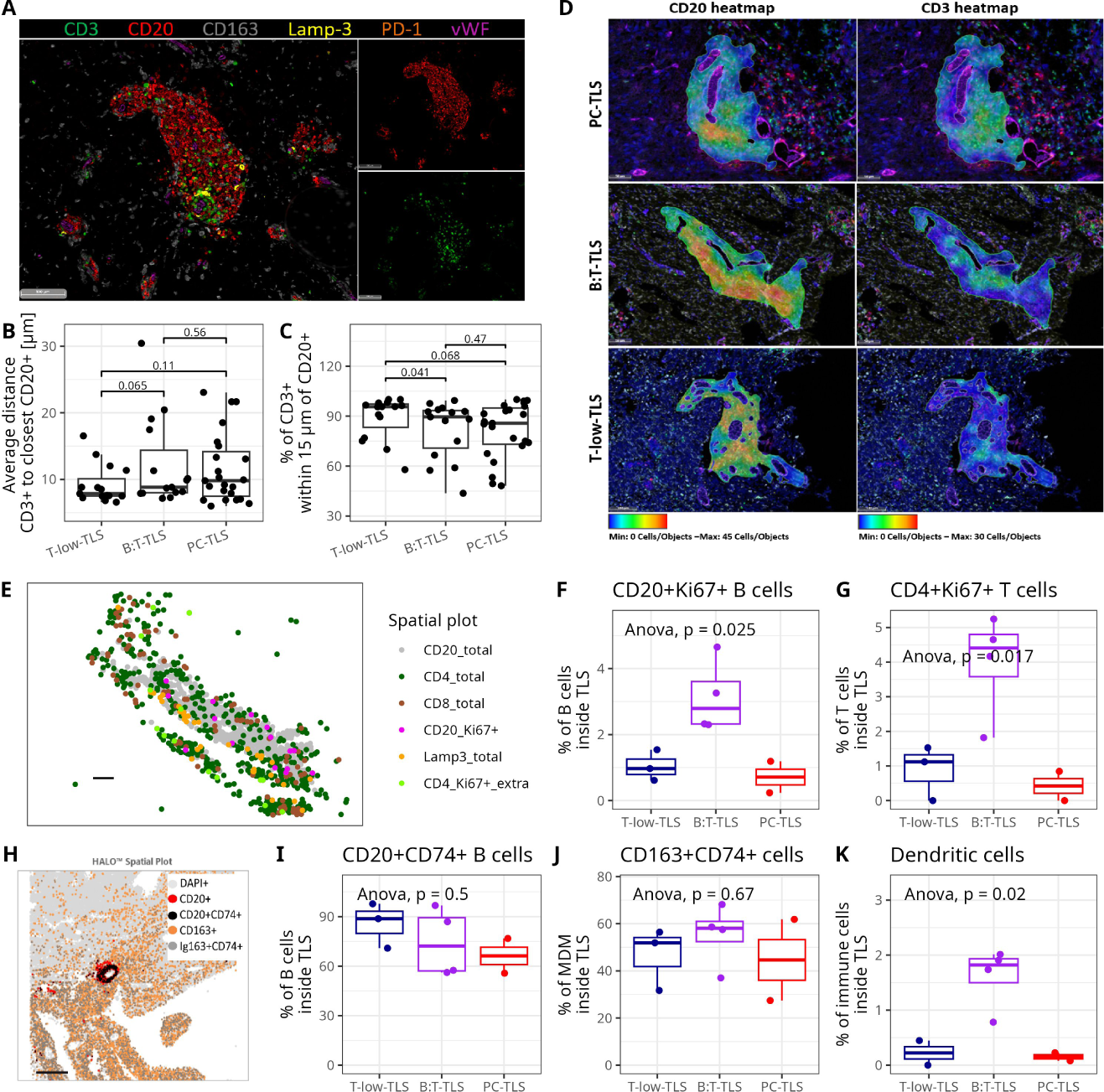
B:T-TLS and PC-TLS show evidence for dynamic adaptive immune responses. **A**: 7-plex immunofluorescence staining highlighting CD3+ T cells and CD20+ B cell localization within a T-low TLS. **B**: Average distance between a CD3+ cell and the closest CD20+ cell for each TLS measured in 7-plex immunofluorescence images. **C**: Percentage of CD3+ cells within 15 µm radius of CD20+ cells. The lower the percentage, the higher the number of CD3+ T cells without immediate CD20+ B cell neighbors. For B and C panels, each dot indicates the measured value for one TLS structure. TLS are grouped according to TLS subtype (T-low-TLS, B:T-TLS, PC-TLS). Boxplots indicate median, 25% and 75% quantiles. P-values in B and C are generated from Wilcoxon test. **D**: Density heatmaps representing the density of CD20+ B cells (left) and CD3+ T cells (right) within different TLS. Color scale represents densities between 0 and 45 B cells/area (25µm radius) and between 0 and 30 T cells/area (25µm radius). Representative examples of each TLS subtype are displayed. **E:** Location of different cell phenotypes in one B:T-TLS, data extracted from 32-plex imaging data. Colors indicate phenotypes: CD20+ B cells (grey), CD4+ T cells (dark green), CD8+ T cells (brown), Ki67+CD20+ B cells (pink), Lamp3+ dendritic cells (orange), Ki67+CD4+ T cells (light green). Scale bar 100 µm. **F+G**: Percentage of CD20+Ki67+ among CD20+ B cells (F) and percentage of CD4+Ki67+ T cells among CD4+ T cells (G) inside TLS areas. Each dot represents the percentage for one TLS. TLS are grouped and colored according to TLS subtypes. Boxplots indicate median, 25% and 75% quantiles. P-values calculated using ANOVA. **H**: Location of different cell phenotypes within a single TLS, plot extracted from 32-plex imaging data. Colors indicate different phenotypes: CD20+ B cells (red), CD20+CD74+ B cells (black), CD163+ macrophages (orange), CD163+CD74+ macrophages (dark grey), other DAPI+ cells (light grey). Scale bar: 500 µm. **I:** Percentage of CD20+CD74+ among B cells inside TLS area. **J:** Percentage of CD163+CD74+ among CD163+ macrophages within the TLS. **I**: Percentage of Lamp3+ dendritic cells among all immune cells identified within the TLS. Colors and statistics in I, J, K as in F.

Highplex immunofluorescence staining identified CD20+Ki67+ and CD4+Ki67+ proliferating B and T cells, which accumulated in the corresponding B cell- and T cell-rich areas of TLS, indicating proliferation and clonal expansion of adaptive immune cells within TLS (**Fig. 4E**). Proliferating (Ki67+) B cells and CD4 T cells were highest in B:T-TLS, comprising ∼3% of all B cells and ∼4% of all CD4 T cells (**Fig. 4F and 4G**). In line with these finding, the maximal B cell density was highest in the B:T-TLS cluster (**Fig. S5A**), suggestive of B cell aggregation by B cell clonal expansion, whereas the average density of B cells was comparable throughout the different TLS types (**Fig. S5B**). Our data further provided evidence for CD4+PD1+ and CD4+PD1+ICOS+ T cells of the follicular helper T cell phenotypes, with no significant differences between the TLS subtypes (**Fig. S5E**). CD4+GranzB+ T cells were enriched inside B:T-TLS (**Fig. S5E**).

Between 60-90% of all B cells and 30-80% of all macrophages within TLS displayed a CD74+ phenotype, indicative of antigen presentation function (**Fig. 4H-4J, S5C and S5I)**. In addition, CD74+ macrophages localized outside the TLS, whereas B cells were scarcely detected outside TLS regions (**Fig. 4H and S5D and S5J**). Lamp3+ dendritic cells localized close to T cells (**Fig. 4E**). They constituted approximately 2% of all immune cells within B:T-TLS and less than 0.5% in the remaining subtypes, indicating highest levels of professional antigen presentation in B:T-TLS compared to the other two TLS subtypes (**Fig. 4K**). We did not find evidence for CD23+ follicular dendritic cells within glioma-associated TLS.

### Plasma cell formation coincides with extracellular matrix remodeling and CD8+ tumor infiltration

We next investigated the cellular and immunological features of plasma cell formation within the TLS. Highplex immunofluorescence staining revealed the presence of class-switched plasma cells with IgG+CD38+MZB1+ and IgA+CD38+MZB1+ phenotypes inside and outside of TLS areas (**Fig. 5A-C**). The highest percentage of plasma cells outside the TLS was in proximity to PC-TLS and some B:T-TLS, suggesting the formation and egression of plasma cells from these TLS (**Fig. 5A-5C**).

**Figure 5:**
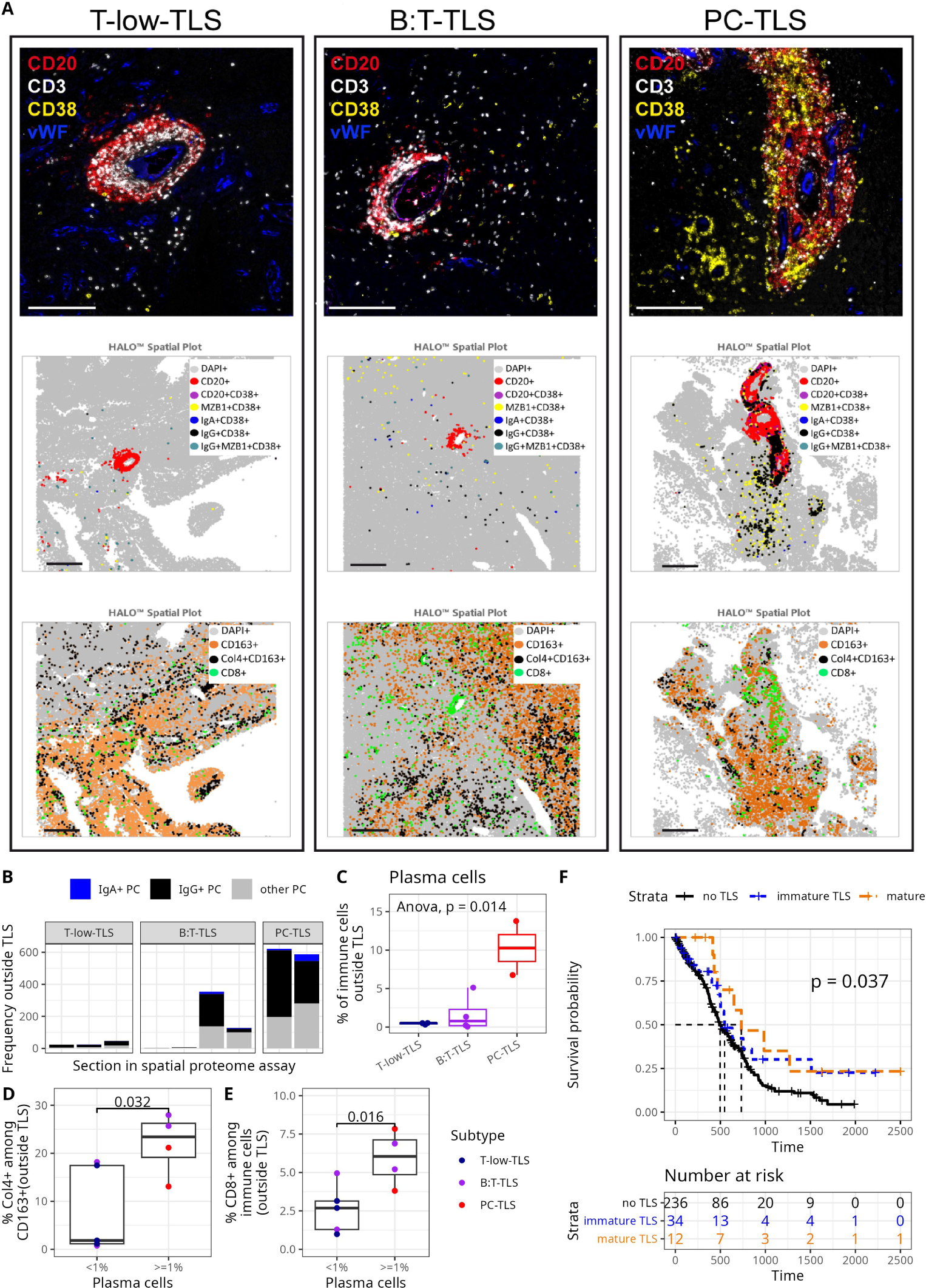
Plasma cell formation by B:T- and PC-TLS is linked to extracellular matrix remodeling and CD8+ tumor infiltration. **A:** Spatial proteomics analysis of plasma cells within the three TLS subtypes. Top panels: Representative composite images from 32-plex immunofluorescence staining with CD20+ B cells (red), CD3+ T cells (white), CD38+ plasma cells (yellow) and vWF+ endothelial cells (blue) within T-low-TLS, B:T-TLS and PC-TLS. Scale bars: 200 µm. Middle panels: Representative plots of the three TLS subtypes within the tumor area of ∼10 mm^2^, displaying spatial distribution of CD20+ B cells (red) and corresponding B- and PC-subtypes: CD20+CD38+ (violet), MZB1+CD38+ (yellow), IgA+CD38+ (blue), IgG+CD38+ (black), IgG+MZB1+CD38+ (cyan). Scale bars: 500 µm. Bottom panels: Spatial distribution plots of representative TLS subtypes, displaying CD8+ T cells (green dots), CD163+ macrophages (orange dots) and CD163+Col4+ macrophages (black dots). Scale bars: 500 µm. **B:** Absolute frequencies of plasma cells outside of the TLS within the ∼10 mm^2^ imaging area. Colors indicate IgA+ plasma cells (blue), IgG+ plasma cells (black) and IgA-IgG-plasma cell (grey). Plasma cells are defined as CD38+MZB1+. **C:** Percentage of plasma cells among all immune cells outside TLS regions. Each dot represents the percentage for one TLS. TLS are grouped and colored according to the TLS subtypes. Boxplots indicate median, 25% and 75% quantiles. P-values from ANOVA. **D:** Percentage of CD163+Col4+ among all CD163+ macrophages outside TLS regions, grouped according to the percentage of plasma cells found outside of TLS (< 1% and >=1% of immune cells). Colors as in C, p-value from Wilcoxon test. **E**: Percentage of CD8+ T cells among all immune cells outside TLS regions, grouped according to the percentage of plasma cells found outside of the TLS (< 1% and >=1% of immune cells). Colors as in C, p-value from Wilcoxon test. **F:** Overall patient survival analysis for patients with IDH-wt glioma without TLS (black), patients with small immune aggregates (< 500 total cells, blue), patients with mature TLS (orange). Percentage of patients with > 4-year overall survival per group: 4% (9/236) for no TLS, 12% (4/34) for small immune aggregates, 17% (2/12) for mature TLS group. p-value indicated in figure from a statistical test to detect an improved overall survival trend over the three groups from no TLS, over small immune aggregate to mature TLS group. P values for individual comparisons: no TLS vs. immature TLS (p = 0.097), no TLS vs. mature TLS (p = 0.051), immature TLS vs. mature TLS (p = 0.5).

When correlating the levels of plasma cells formed by TLS to other characteristics of the TME, we observed that plasma cell formation coincided with overall higher immune cell infiltration and extracellular matrix remodeling. For this, we divided the tumors into two groups dependent on whether plasma cells constituted less than 1% or more than 1% of immune cells outside of TLS. Macrophages in tumors with high plasma cell levels were more likely to exhibit a CD163+Col4+ phenotype than macrophages in tumors with low plasma cell levels (**Fig. 5D; Fig. S3**). Similarly, the frequency of CD8+ T cell outside the TLS significantly increased in the presence of plasma cells (**Fig. 5E; Fig. S3**). Overall survival of patients was improved with increasing TLS maturation quality from tumors with no TLS over tumors harboring smaller immune aggregates (defined as TLS with a total cell number < 500) to tumors enriched in B:T-or PC-TLS (mature TLS) (**Fig. 5F**). This finding supports the notion that mature TLS are immunological more functional and thus contribute to prolonged patient survival.

### B:T-TLS, PC-TLS and T-low-TLS represent independent differentiation states of TLS formation

Molecular spatial profiling suggests that the three TLS subtypes represent qualitatively distinct immunological states of TLS. We grouped the 60 TLS profiled in spatial transcriptomic according to their tumor of origin, and thereby determined the number of TLS per TLS subtype in each tumor (in total 20 tumors). We found that although different TLS subtypes could be observed within one tumor, usually one TLS type was overrepresented (**Fig. 6A**). This allowed us to group the tumors into 2 groups: *mature-TLS tumors*, which contained a majority of B:T- and PC-TLS, and *T-low-TLS tumors*, which were enriched in T-low-TLS (**Fig. 6A**). Given the under-representation of T cells within T-low-TLS, we hypothesize that these structures failed to reach a mature germinal center-like state due to insufficient early T cell recruitment into the perivascular space. We therefore quantified the percentages of B and T cells in perivascular immune aggregates of all sizes within T-low-TLS tumors and compared them to mature TLS tumors (**Fig. 6B**). Small perivascular aggregates (< 100 cells), which presumably constitute the early stages of TLS formation, had low T cell frequencies and almost exclusively consisted of B cells in T-low-TLS tumors (**Fig. 6B and 6C**). This observation suggests that T-low-TLS tumors lacked the small T cell-enriched perivascular aggregates that could be observed in mature TLS tumors (**Fig. 6B and 6C**).

**Figure 6:**
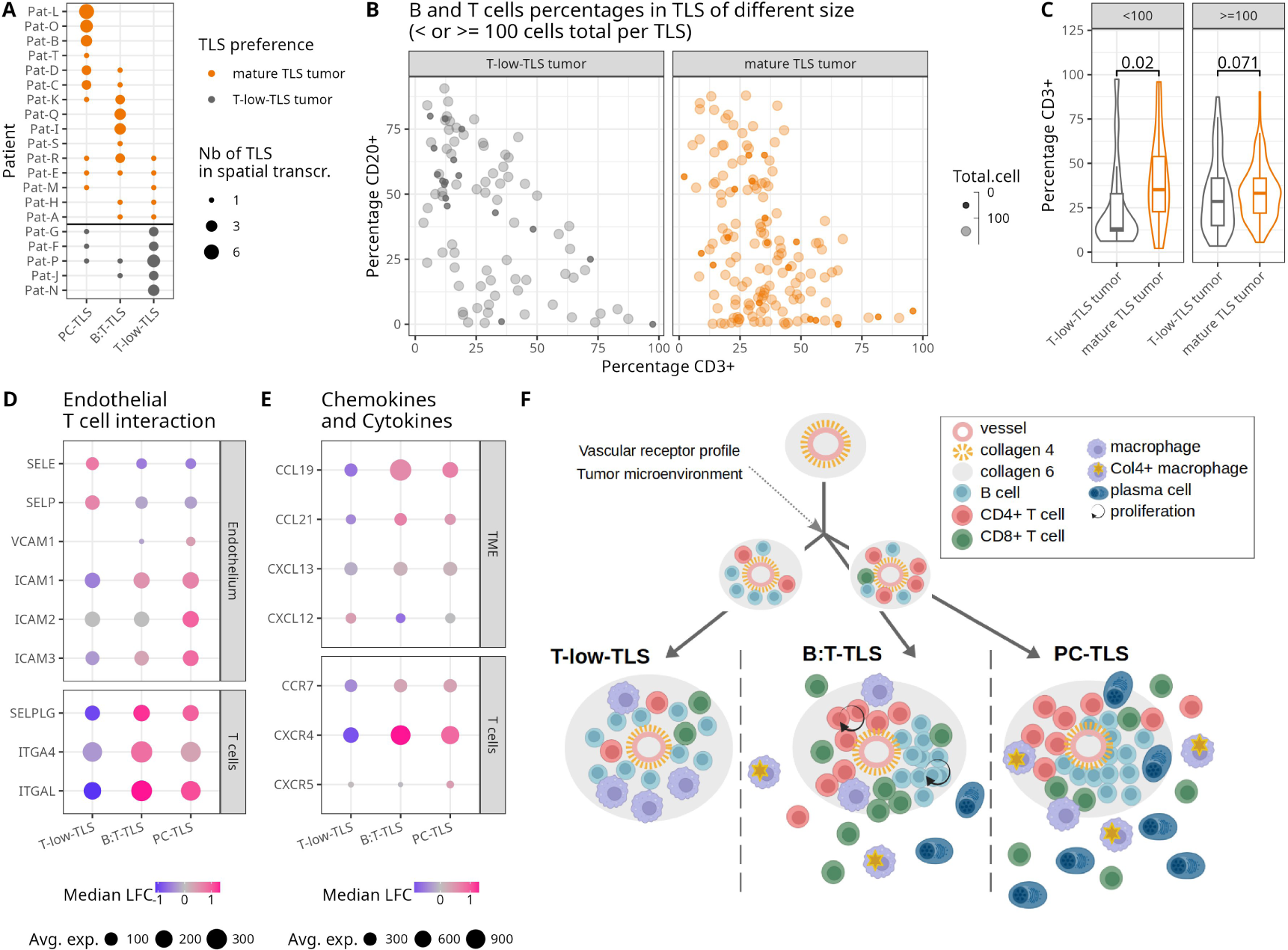
B:T-TLS, PC-TLS and T-low-TLS represent independent differentiation states of TLS formation. A: Number of TLS from each TLS subtype (x-axis) identified in each tumor (y-axis). Size indicates number of TLS per TLS subtype and tumor (from spatial transcriptomics data). Color indicates whether the percentage of T-low-TLS in each tumor was >50% (grey, T-low-TLS tumors) or <=50% (orange, mature-TLS tumors). **B**: B cell percentage (y-axis) and T cell percentage (x-axis) in immue aggregates with a total cell number <100 (small dots) and >=100 (larger dots). T-low-TLS tumor are shown on the left in gray, and mature-TLS tumor on the right in orange. The figure shows data from 20 tumors (whole-slide sections analyzed using 7-plex immunofluorescence staining). **C**: Percentage of CD3+ T cells in TLS with a total cell number <100 (left) and >=100 (right). Color as indicated in A. **D and E**: Expression profile of genes mediating the interaction of endothelial cells and T cells (D) and chemokine/cytokine signaling from the TME to T cells (E) during transendothelial migration. Regions of interest were grouped into different TLS subtypes. Dot size indicates the average normalized expression of a gene across TLS of a specific subtype. Color indicates median logarithmic fold change of gene expression in a TLS subgroup compared to the overall mean across all TLS. Data for D and E were obtained from spatial transcriptome profiling. **F**: Model of TLS differentiation into the three different TLS subtypes (T-low-TLS, B:T-TLS and PC-TLS). The maturation into B:T- and PC-TLS is facilitated by an efficient infiltration of T cells into the early perivascular immune aggregates, which is driven by vascular receptor profile and TME.

Since T cell migration across the BBB is a multistep process orchestrated by a plethora of adhesion receptors and signaling molecules^36^, we next investigated the expression of transcripts of ligands and receptors involved in T lymphocyte extravasation into the brain tumor parenchyma (**Fig. 6D**). While T-low-TLS transcriptional profiles, in comparison to the other two TLS subtypes, displayed higher levels of receptors involved in the initial steps of the adhesion cascade (*SELE*, *SELP*; **Fig. 6D**), they showed lower levels of vascular adhesion receptors and chemokines necessary for firm attachment and subsequent diapedesis (in particular *ICAM1*, *ICAM2*, *ICAM3*, *CCL19, CCL21*; **Fig. 6D and 6E**). The lower expression of these molecules could explain the lower T cell frequencies observed in the perivascular niche of T-low-TLS tumors.

These finding supports the hypothesis that the immunological states defined in the three TLS subtypes represent distinct final differentiation states rather than temporal evolutionary stages. In this scenario, the probability of differentiating into one of the TLS subtypes depends on tumor-intrinsic factors (e.g. vascular phenotype, tumor metabolism). This leads to the majority of TLS within one tumor differentiating into the TLS subtype with the highest tumor-specific generation probability (**Fig. 6F**). Indeed, we were able to identify metabolic differences between T-low-TLS tumors and mature-TLS tumors that could explain their TLS preference. Pathways related to cell cycle, glycolysis, hypoxia and VEGF signaling were downregulated in T-low-TLS tumors compared to mature-TLS tumors and TLS-tumors, suggesting that metabolism driven signaling pathways rather than immune cell intrinsic differentiation leads to different TLS subtypes (**Fig. S6A-S6D**). The lower proliferation rate of T-low-TLS tumors compared to mature-TLS tumors was also confirmed when comparing the routine diagnostic Ki67+ score (**Fig. S6E**). In summary, we propose the following model for glioma-associated TLS formation: Small perivasular immune aggregates can differentiate into the three different TLS subtypes (T-low-TLS, B:T-TLS and PC-TLS). The maturation into B:T- and PC-TLS is facilitated by an efficient infiltration of T cells into the early perivascular space (**Fig. 6F**).

## Discussion

In this study, we demonstrate 15% of adult-type diffuse gliomas harbor TLS, which differentiate into mature stages characterized by sustained adaptive immune responses associated with improved patient survival. The incidence of TLS in glioma is lower compared to other solid cancers such as pancreatic, rectal, lung and ovarian cancer^19,37–39^. Nevertheless, our study challenges the paradigm of glioma being a *bona fide* immune-cold tumor, since it demonstrates the existence and clinical relevance of dynamic adaptive immune responses. Importantly, we show that patient stratification according to TLS quality has a higher prognostic value than stratification simply based on the presence of B cell aggregates. Our work delineates molecular and cellular correlates of TLS formation and differentiation in glioma, and thereby contributes to greater understanding of adaptive anti-glioma immunity.

In many solid tumors, the availability of mutated tumor-specific antigens due to high TMB values is associated with improved adaptive immune responses^30^. In glioma however, we find that TLS can be formed despite low levels of TMB, highlighting the fact that antigen availability is not the only determinant of adaptive immunity. This is in line with a lack of association between TMB and immunotherapy response in gliomas^40^.

Our comparison of TLS+ and TLS-gliomas indicates a decreased neural tumor signature in TLS+ tumors. This is in line with a recent study comparing brain tumors with high- and low-neural signature. This study found increased ECM and immune cell signatures in gliomas with low-neural signature, which were also associated with improved overall and progression-free survival^41^. Furthermore, our study demonstrates non-canonical collagen expression by stromal components of the TME and increased perivascular ECM remodeling in TLS+ tumors that could be a prerequisite for TLS formation. This non-canonical collagen expression (Col4 low/Col6 high) is observed at the earliest stage of TLS formation and presents before immune cells accumulate within the perivascular niche. Supporting our hypothesis, constitutive upregulation of Col6 in the perivascular niche of high-grade glioma but not normal brain has been reported^32,33^. The ECM remodeling in TLS+ tumors is accompanied by expression of ECM-remodeling enzymes and internalization of collagen components by tumor-associated macrophages. Remodeling of the blood-brain barrier extracellular matrix components by matrix-metalloproteases has previously been linked to blood-brain barrier opening^42^, which could improve immune cell infiltration.

Our study further demonstrates the clinical relevance and delineates the cellular and molecular architecture of three qualitatively distinct TLS differentiation states identified in the TLS+ adult-type diffuse gliomas. Two of the intra-tumoral TLS subtypes identified in our study, namely B:T-TLS and PC-TLS, elicit features of ongoing adaptive immune responses. Despite the absence of CD23+ follicular dendritic cells, these functional TLS are capable of producing IgA+ and IgG+ class-switched plasma cells that egress from the TLS area, in line with previous reports on tumor infiltrating plasma cells in ovarian cancer^43^ and renal cell cancer^25^. The B:T-TLS subtype features Lamp3+ DCs that predominantly localize to the T cell zone and therefore shows the strongest similarity to mature TLS described in other tumor entities, such as lung tumors^38^. The increased tumor infiltration by CD8+ T cells around mature TLS is comparable to the findings of previous studies in melanoma, sarcoma, ovarian and lung cancer, where mature TLS and presence of dendritic cells (Lamp3+) were associated with tumor-infiltrating CD8+ T cells^15,16,37,38^.

A smaller subset of glioma-associated TLS (T-low-TLS subtype) is characterized by a less-functional phenotype with high levels of CD163+ macrophages and low levels of T cells and is therefore evocative of previously described lympho-myeloid aggregates^20^. In our study, these structures have markedly lower plasma cell forming capacity than B:T- and PC-TLS and are associated with lower levels of T cell infiltration into the tumor parenchyma. T-low-TLS tumors lack T cell rich perivascular niches and show a reduced expression of molecules necessary for T cell trafficking across the BBB, suggesting that the endothelium in these tumors is less attractive to T cells. Indeed, others have positively correlated T cell, in particular SELL+ T cell, abundance in TLS with the expression of T cell capture molecules in lung cancer^44^. T-low-TLS tumors do not only seem to attract fewer T cells, but also showed significantly reduced levels of VEGF signaling, glycolysis and other hypoxia-related pathways. These findings are in line with reports that hypoxia-induced expression of HIF-1a and subsequent metabolic reprogramming regulates T cell expansion and differentiation^45,46^. As such, our findings support the notion that local microenvironmental signals drive TLS expansion and maturation in gliomas and further underscore the pivotal role of the glioma microenvironment in shaping the tumor specific immunology landscape. Of note, T-low-TLS were found in both IDH-wt and IDH-mut gliomas, suggesting that the metabolic mechanisms involved in their formation are somewhat independent of the IDH-mut specific metabolite R-2-hydroxyglutarate that has previously been associated with dysfunction of macrophages and dendritic cells^47,48^.

In summary our multi-modal study comprehensively assesses the clinical relevance of TLS and identifies the cellular mechanisms that promote TLS maturation in adult-type diffuse glioma. The relevance of pathway-based glioblastoma classification as opposed to lineage-based classifications has only recently been emphasized (Garofano et al., Nature Cancer 2021). Our multi-modal approach revealed a previously unknown type of glioblastoma which is informative of patient survival, characterized by the presence of mature TLS that can be identified in pathology tissue sections. These insights are of potential relevance for a redefined patient stratification in glioblastoma immunotherapy trials, based on the presence or absence of mature TLS, as well as for the development of targeted therapies for induction or maturation of TLS.

## Acknowledgments

The authors would like to thank Samira Ortega Iannazzo, Adrien Jolly and Stefan Liebner for helpful feedback on the manuscript as well as Marie Cromm for facilitating metadata annotation. Furthermore, the authors thank the University Cancer Center tumor documentation team (Kristina Goetze, Sandra V. Klein, Andrea Wolf), for providing clinical data and advice for clinical data analysis. Stella Breuer participated in the neuroradiological assessment. J.H.L. is supported by the Uniscientia foundation. P.C. was supported by the Edinger foundation. L.M.H and K.I. are supported by funding from the Mildred Scheel Career Center Frankfurt (Deutsche Krebshilfe). K.I. and A.S. are funded by the Frankfurt Cancer Institute (LOEWE program). K.H.P. received funding from the Clinical translational program “Individualized immunotherapy for gliomas” (Frankfurt Cancer Institute, LOEWE program). The Immunomonitoring Platform (K.H.P.) is supported by Frankfurt Cancer Institute (LOEWE program), German Consortium for Translational Cancer Research (DKTK), Uniscientia foundation and Edinger foundation. Biorender was used to create figure 1A, 3A, and 6F.

## Author contributions

P.C., J.H.L., T.S., J.S., J.M. conducted experiments. A.S., L.M.H., M.K., K.I. implemented software and curated data. P.C., J.H.L., A.S., J.M., K.I. performed data analysis. E.H., E.S., M.C.B. provided clinical data. P.C., J.H.L., A.S., J.M., K.H.P., Y.R., K.I. wrote the paper. K.I., Y.R. and K.H.P. designed the study and developed methodology.

## Resource availability

### Lead contact

Katharina Imkeller

### Materials availability

This study did not generate new unique reagents.

### Data and code availability

Gene expression data from bulk RNA sequencing and related metadata is available from GEO under the accession number GENEEXPRESSIONGEO (will be made publicly available after peer review). Gene expression data from GeoMx DSP spatial profiling and related metadata is available from GEO under the accession number GEOMXGEO (will be made publicly available after peer review). Softwares and codes used for data collection and analyses were outlined in the methods. Any additional information required to reanalyze the data reported in this paper will be made available from the lead contact upon reasonable request.

## SUPPLEMENTARY FIGURES

**Fig. S1:**
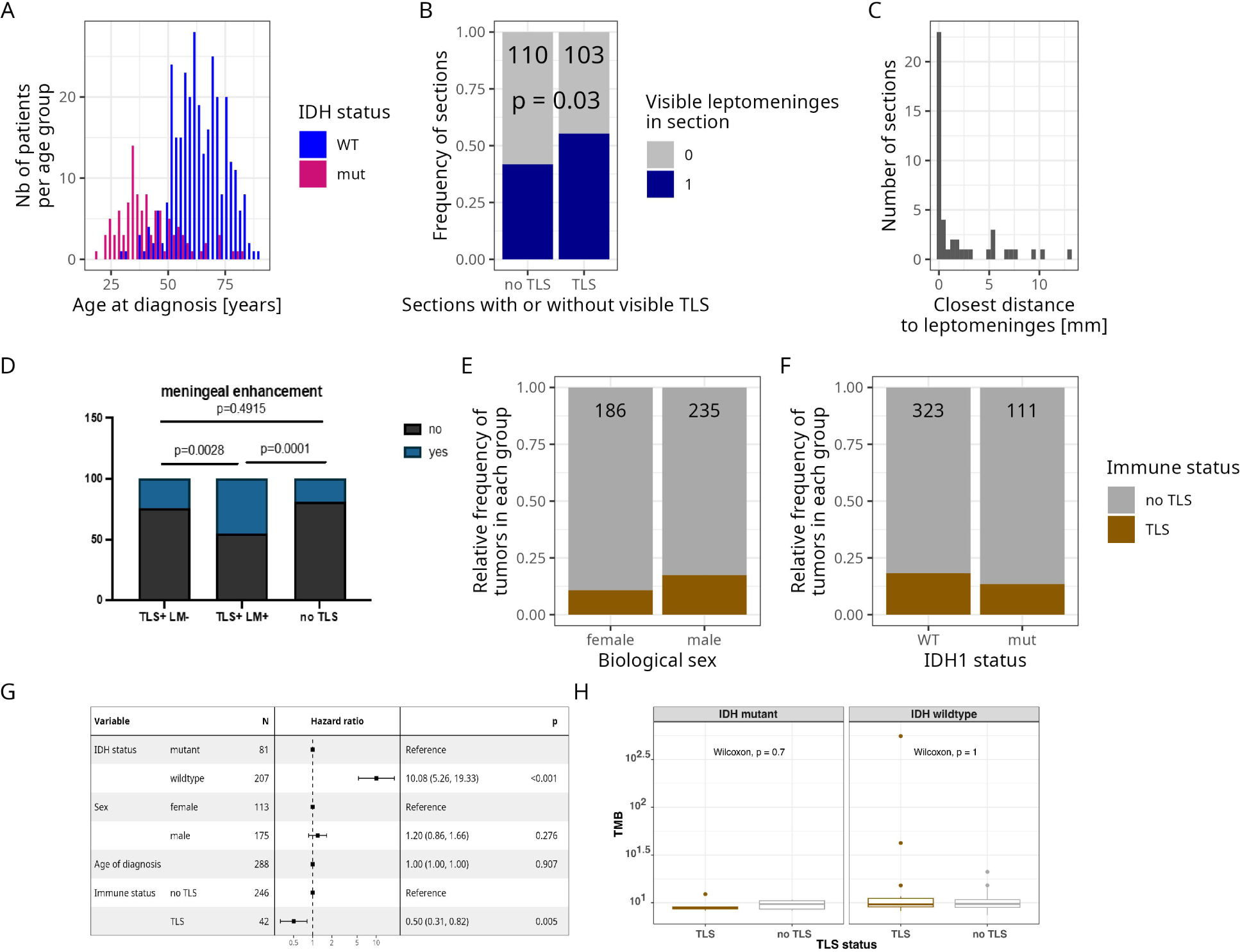
Clinical features of TLS+ gliomas and association with leptomeninges. **A**: Histogram of the age at diagnosis of patients within the study cohort. Patients with IDH-wt tumors are depicted in blue and patients with IDH-mut tumors in magenta. **B**: Frequency of sections with visible leptomeninges (blue) in TLS+ (n=110) and TLS-(n=103) sections. P-value from Fisher exact test. **C**: Histogram of the closest distance between TLS and the leptomeninges in each of the TLS+ sections analyzed in B. A distance of 0 indicates TLS found within the leptomeninges. **D**: Rates of meningeal enhancement in three tumor groups: tumors with at least one TLS in leptomeninges (TLS+ LM+, n=11), TLS+ tumors without TLS in leptomeninges (TLS+ LM-, n=41), and TLS-tumors (n=58). P-values calculated by Fisher’s exact test. **E and F**: Relative frequency of TLS+ (brown) and TLS-(gray) tumors identified in D) female (n=186) vs. male (n=235) and E) IDH-wt (n=323) vs. IDH-mut (n=111) tumors. According to the multinomial logistic regression, TLS presence is higher in male patients (p = 0.02), but independent of the IDH status (p = 0.59). **G**: A forest plot depicting the hazard ratio (HR) and 95% CI of the association between tumor progression and different clinical parameters (tumor IDH mutational status, patient sex, patient age at diagnosis, TLS+/TLS-status). We calculated the number of days to reach progression after diagnosis based on clinical follow-up data. For the patients whom no progression was detected up until their last visit, their data was right censored. A multinomial Cox proportinal hazard regression model was applied to determine statistical significance. **H**: Boxplots displaying the tumor mutational burden (TMB) value for TLS+ (brown) and TLS-(gray) in IDH-mut and IDH-wt gliomas.

**Fig. S2:**
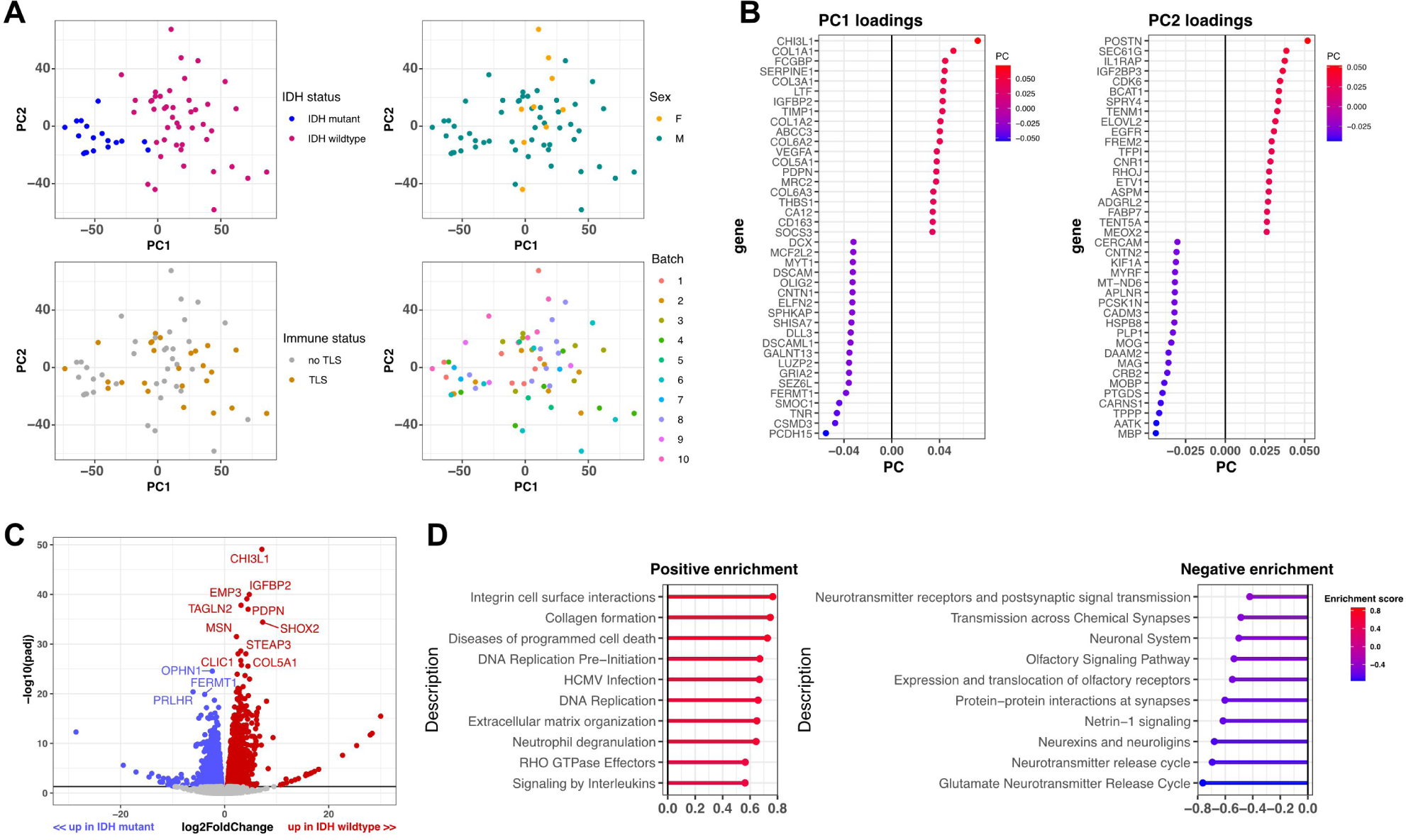
Principal component analysis (PCA) of bulk RNA sequencing data. **A**: Principal component analysis (PCA) of bulk RNA sequencing transcriptional profiles of TLS+ and TLS-gliomas. Different PCA plots show the distribution of IDH mutational status, patient biological sex, TLS status and batch (i.e. experimental runs). **B**: Loadings of the principal components PC1 and PC2. The 15 genes with the highest and lowest loading coefficients are displayed. **C**: Volcano plot displaying differentially expressed genes between IDH wildtype and IDH mutant glioma. Labeled genes had a >2-fold change and adjusted p value <0.05. **D**: GSEA results. Pathway enriched in IDH wildtype glioma are shown in red, those enriched in IDH mutant glioma are shown in blue.

**Fig. S3:**
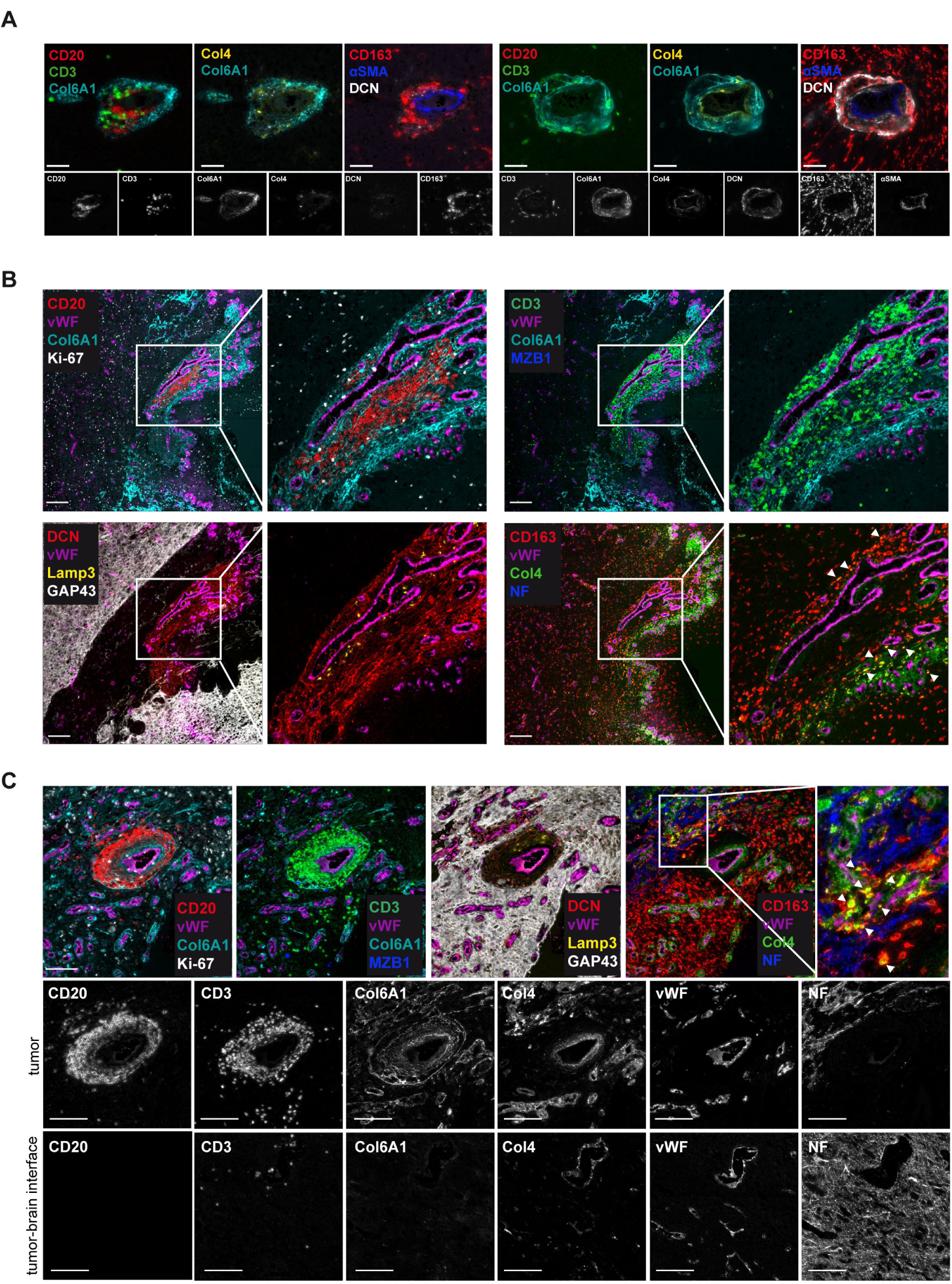
Non-canonical collagen expression within the perivascular niche of glioma TLS. **A**: Representative immunofluorescence images (COMET, extracted from 32-plex panel) and identification of CD20+ B-cells (red) and CD3+ T-cells (green) in the perivascular niche: Col6A1 (cyan), Col4+ (yellow), DCN (white), αSMA+ cells (blue), CD163+ myeloid cells (red), and corresponding single channel images. Scale bars: 50 µm. **B**: Example of a TLS embedded within dense Col6A1 network (cyan) surrounding vWF (violet) glioma vessel. Upper panel: CD20+ B-cell (red, left) and CD3+ T-cell (green, right) aggregation within Col6A1 network (cyan), highlighted in boxed areas at higher magnification. Ki67 (white), MZB1 (blue). Lower panel: Lamp3+ dendritic (yellow, left), CD163+ (red) and Col4+ (green) cell type identification (right) with boxed insets at higher magnification. Arrows highlighting CD163+Col4+ positive macrophages. Absence of NF (blue) as a sign for intra-tumoral TLS localization. DCN (red), GAP43 (white). Scale bars: 200 µm. **C**: Upper panel: Composite immunofluorescence images of a TLS within a less densed Col6A1+ (cyan) perivascular ECM network (compare to B). vWF (violet), MZB1 plasma cells (blue), Lamp-3+ (yellow). Arrows: CD163+ macrophage area (red) with Col4+ fragments (green). Scale bar: 200 µm. Lower panels displaying single-plex images of the same ROIs, and comparison to non-tumor area. Marker display: CD20, CD3, Col6A1, Col4, vWF and NF (demarcation of the brain-tumor interface). Col6A1 is absent in non-tumor region. Scale bars: 200 µm.

**Fig. S4:**
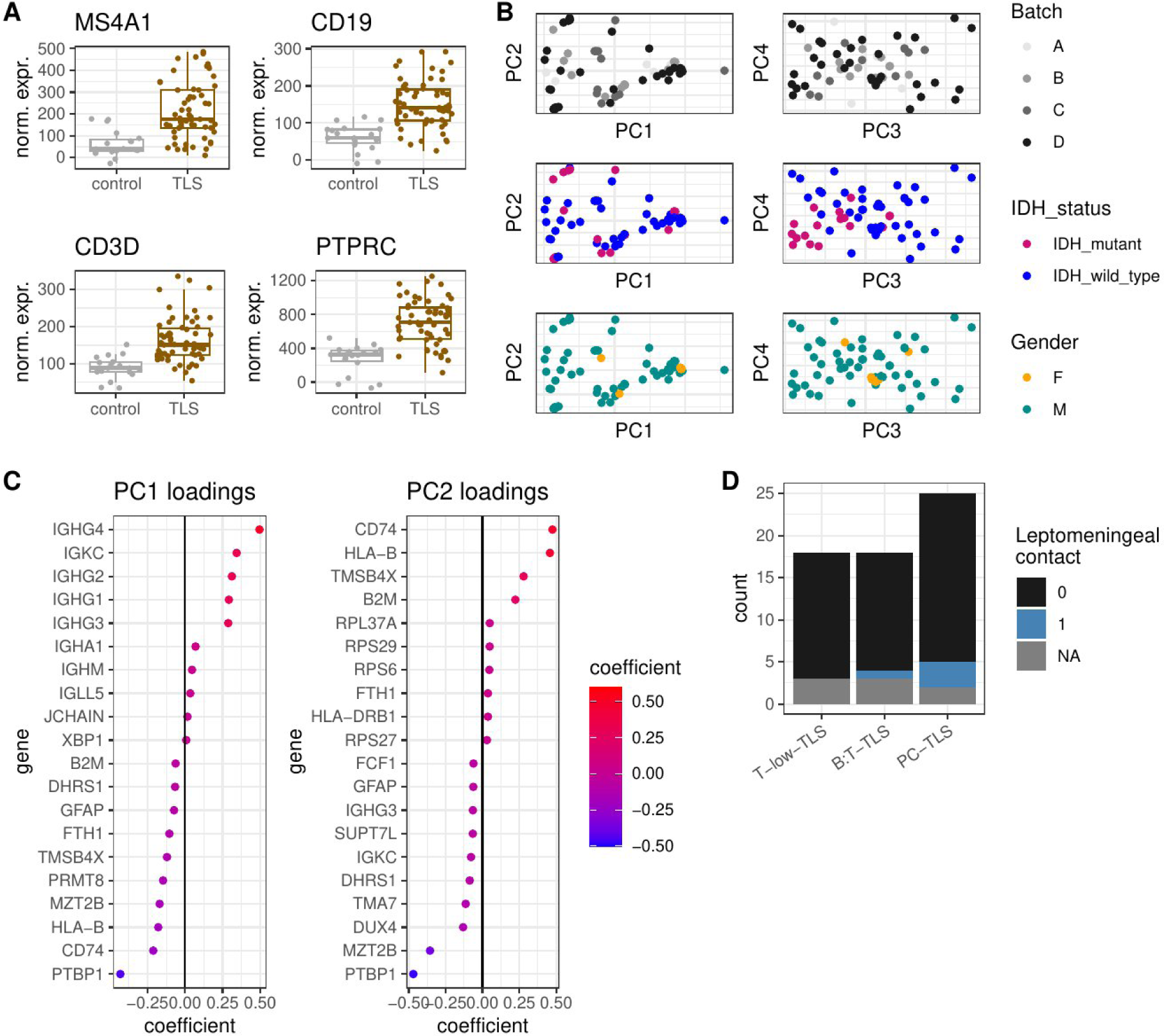
Transcriptional profiles of TLS across patients and locations. **A**: Normalized expression levels of MS4A1 (encoding CD20), CD19, CD3D and PTPRC in tumor rich control regions (gray) and CD20+ TLS regions (brown). Data from spatial transcriptomics. **B**: Principal component analysis of transcriptional profiles of CD20+ TLS regions of interest. PC1-PC4 are displayed and TLS colored according to experimental batch (first row), IDH mutational status (second row) and biological sex (third row). Percentage of variance explained by the principal components: PC1 - 42%, PC2 - 20%, PC3 - 7%, PC4 - 5%. **C**: Loadings of principal components 1-2. The 10 genes with highest and lowest loading coefficient are shown for each PC. **D**: Frequency of TLS without (black) or with (blue) leptomeningeal contact in each of the TLS subtypes. TLS for which leptomeningeal contact could not be determined are shown in gray.

**Fig. S5:**
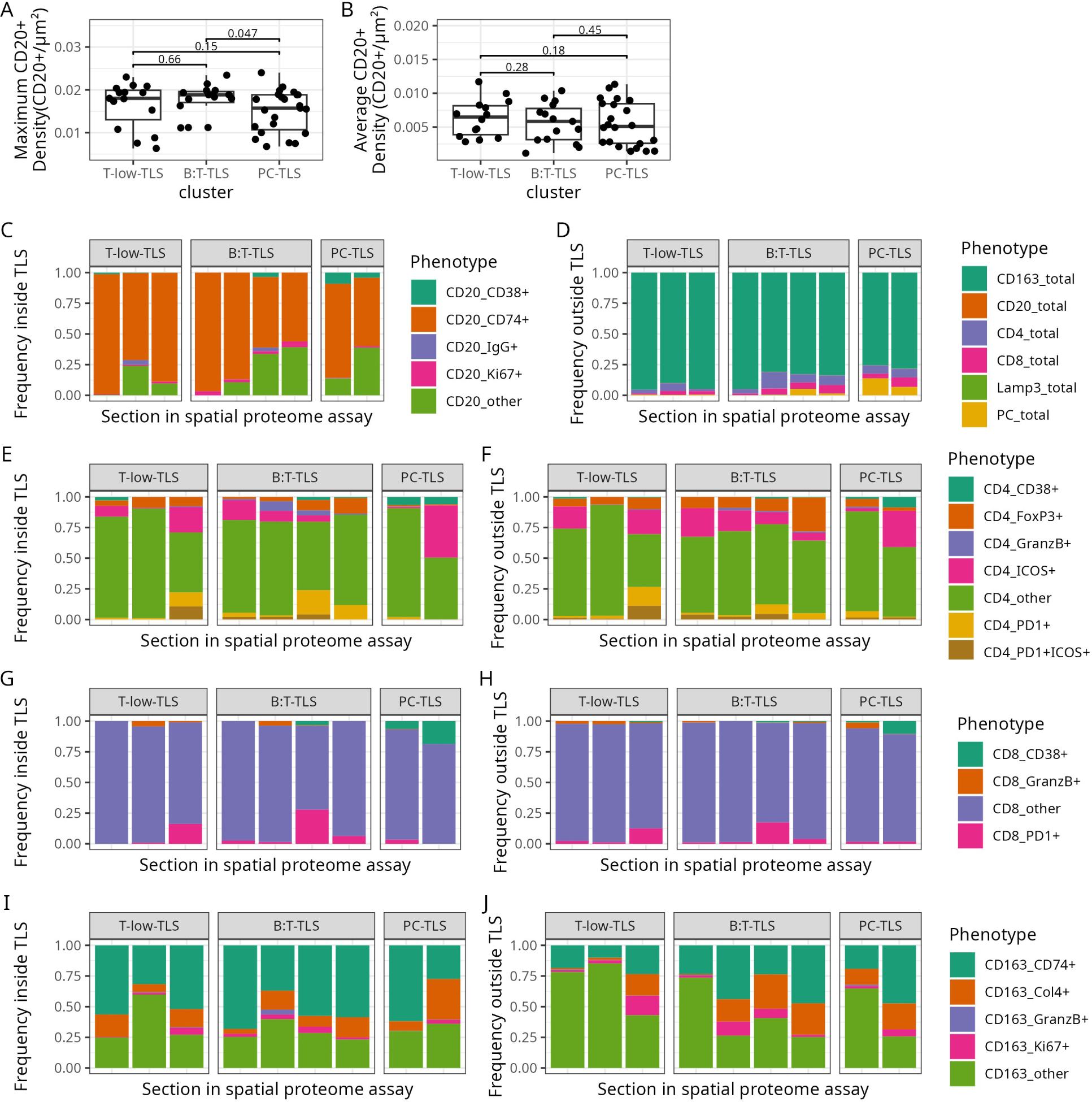
Multimodal imaging approach. **A**: Maximal CD20+ B cell density in TLS from different subtypes. **B**: Average CD20+ B cell density in TLS from different subtypes. Units for A and B are CD20+ cells/µm^2^. Data for A and B from 7-plex immunofluorescence images. **C**: Relative frequencies of B cell phenotypes detected within the TLS, separated by subtypes. **D**: Relative frequencies of immune cell types detected outside of the TLS, grouped by subtypes. **E and F**: Relative frequencies of CD4+ T cell phenotypes detected within (E) and outside of (F) the TLS, grouped by subtypes. **G and H**: Relative frequencies of CD8+ T cell phenotypes detected within (G) and outside of (H) the TLS, grouped by subtypes. **I and J**: Relative frequencies of CD163+ macrophage phenotypes detected within (I) and outside (J) of the TLS, grouped by subtypes. C-J: Data from highplex immunofluorescence staining.

**Fig. S6:**
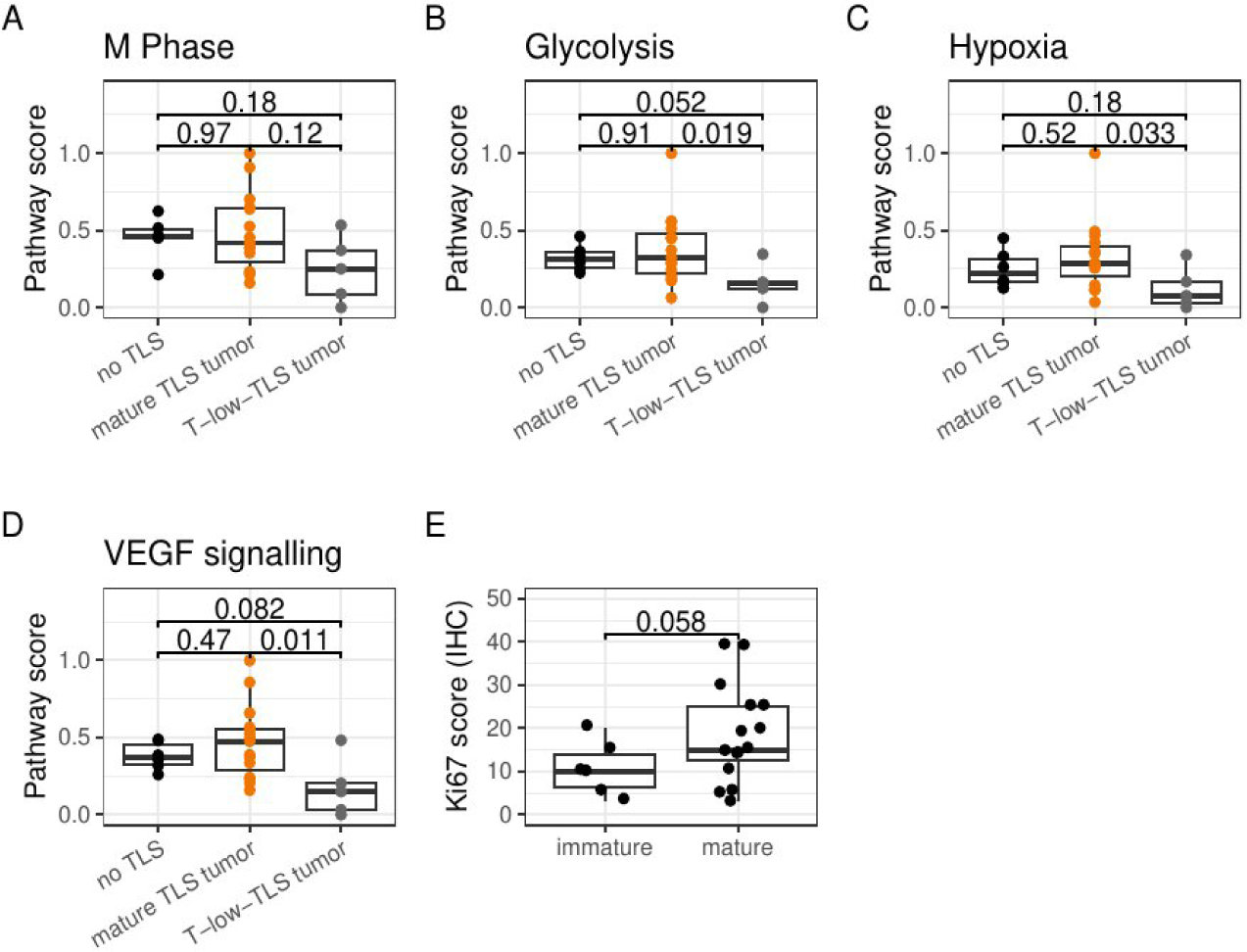
Differences in tumor metabolism between T-low-TLS tumors and mature-TLS tumors. **A-D**: Pathway score for Reactome pathways M-Phase (A; R-HSA-68886), Glycolysis (B; R-HSA-70171), Cellular response to hypoxia (C; R-HSA-1234174) and signaling by VEGF (D; R-HSA-194138) in gliomas without TLS (black), mature-TLS tumors (orange) and T-low-TLS tumors (gray). **B**: Ki67 score in mature-TLS tumors and T-low-TLS tumors. P-value in all panels from Wilcoxon test.

## SUPPLEMENTARY TABLES

**Supplementary Table 1**: Results from differential gene expression analysis comparing TLS+ (n=44) and TLS-(n=20) adult-type diffuse glioma tumors. Gene list only contains genes with p adjusted value (padj) of less than 0.05.

**Supplementary Table 2**: Marker combinations used to define cell phenotypes in 32-plex immunofluorescence data. This table indicates which marker combinations were used to define cellular phenotypes in 32-plex immunofluorescence data. Each row represents one cellular phenotype – the phenotype name is indicated in the first column “phenotype”. The other columns indicate the usage of specific markers for phenotype definition: “1” = cells need to be positive for this marker, “-1” = cells need to be negative for this marker in order to be labeled with the corresponding phenotype. “0” = the marker was not used to define the phenotype. One segmented cell object can have multiple phenotypes. For each immune cell population (CD4+ T cells, CD20+ B cells, plasma cells, CD8+ T cells, CD163+ macrophages and Lamp3+ dendritic cells) the phenotype with suffix “total” comprises all cells of the population. The phenotypes are defined such, that the number of cells in the “total” population corresponds exactly to the sum of cells in the subpopulations. For plasma cells for example: PC_total = (PC_CD38+MZB1+IgG+) + (PC_CD38+MZB1+IgA+) + (PC_other). The subpopulations with suffix “_extra” represent more specific marker profiles and overlap with other subpopulations.

**Supplementary Table 3:** List of antibodies

**Supplementary Table 4:** Clinical information for the sections analyzed in the study. The table presents clinical data for the 642 tumor specimens analyzed, providing detailed information on each specimen and patient. It includes the patient’s sex, whether the specimen was derived from primary or recurrent tumor, tumor localization, TLS classification (TLS or non-TLS), IDH status, methylation class, methylation subclass, age at diagnosis (in days), and survival data. Survival data is specified in days and includes the patient’s survival status (0 = censored, 1 = deceased). For survival calculations, if the survival status is 1, the time difference between diagnosis and death is used; if the survival status is 0, the time difference between diagnosis and the last visit is used. Similarly, for progression-free survival, if progression is detected (status = 1), the time difference between diagnosis and progression is calculated; otherwise, the time to the last visit is calculated. Since some patients have both primary and recurrent sections, clinical data may appear twice for these individuals. Overall survival and progression-free survival analyzes were conducted using primary sections only.

## Methods

### Sample collection and ethical approval

Formalin-fixed paraffin-embedded (FFPE) glioma tissue specimen were collected from the University Cancer Center (UCT) BioBank Frankfurt, Germany and classified according to WHO criteria by board-certified neuropathologists (K.H.P and P.C.). In 316 patients histopathological tumor assessment was accompanied by genome-wide DNA methylation analyzes^49^. Clinical data and tumor samples were obtained upon informed consent of the patients. The study was conducted in accordance with ethical standards such as the Declaration of Helsinki. The study protocol was endorsed by the local ethical committee (SNO-4-2022).

### Patient selection

In an unbiased retrospective approach, we investigated adult-type diffuse IDH-wildtype and IDH-mutant gliomas deposited in the UCT BioBank from 2015 to 2022. Exclusion criteria were patients under the age of 18 years, missing FFPE-tissue blocks and stereotactic biopsies. A total of 642 adult-type diffuse glioma tissue specimens were analyzed, comprising IDH-wildtype (n=452) and IDH-mutant (n=190) GBMs. See also Fig. 1A.

### Clinical data and analysis

Clinical data were obtained through retrospective review of medical records, including radiological records, surgical notes and pathology reports, as well as the UCT Tumor Documentation. Tumor location was assigned based on preoperative imaging studies and intraoperative findings. Tumor recurrence was defined based on histological evidence from subsequent biopsies, with the criterion that the interval between the first and second biopsy was at least three months. IDH R132H IHC (clone H09, 1:50, Dianova, Germany) results were used to determine the presence of IDH mutation. Patients aged 55 years and older were classified as IDH wildtype if IHC staining was negative. In the case of IDH R132H IHC negativity in patients younger than 55 years, we considered the results of methylome analysis which was used to detect rarer IDH mutations based on WHO criteria^4^. The results of methylation subclassification of gliomas were accessed using the platform (v11b4). This classifier employs machine learning algorithms trained on a comprehensive database of DNA methylation profiles from glioma specimens, allowing classification into methylation subclasses^49^. Proximity to meninges was determined by the visibility of meninges in histopathology (n=103 TLS+, n=110 TLS-), and on the presence of meningeal enhancement in post-contrast T1-weighted MRI sequences (n=76 TLS+, n=50 TLS-). Overall survival analysis was performed using the R *survival* package^50^ functions for multinomial cox proportional hazards regression and Kaplan-Maier curves from the *survminer* package. Survival was right censored and the day of last clinical follow up was used for patients that were alive or had and unknown day of death.

### Immunohistochemistry and screening for B cell aggregation

FFPE tissue blocks of 642 glioma specimens were collected and classified according to WHO criteria by board-certified neuropathologists. A representative block of each biopsy was selected for immunohistochemistry staining. Three µm tissue sections were cut (Leica SM2000R microtome, Germany) and mounted onto glass slides (Superfrost Plus, ThermoFisher Scientific, Germany) for IHC staining. IHC staining against CD20 antigen (clone L26, 1:500 dilution; DAKO, Denmark) was performed using standard protocols for the Leica SM2000R automated stainer. For B cell aggregation assessment, CD20-stained glioma specimens were screened using light microscopy and whole slide image analysis. Biopsies with sparse and non-clustered B cells were interpreted as no aggregation, while those containing clusters of CD20+ cells were considered as immune aggregates. Biopsies containing B cell aggregates were then scanned using the Vectra Polaris (Akoya Biosciences, United States) slide scanner and digital image analysis was performed using HALO software (Indica Labs, United States). Immune cell aggregates consisting of ≥ 50 B cells were classified as TLS (according to ^26,51^).

### Multiplex immunofluorescence staining and digital image analysis

Sections of gliomas comprising B cell aggregates of at least 50 cells (n=33) were stained using the Opal Polaris 7-color manual detection kit (#NEL861001KT, Akoya Biosciences), based on the tyramide signal amplification immunostaining technique. Multiplex immunofluorescence staining was performed on a LabSat® research automated staining device (Lunaphore Technologies SA, Switzerland), with an antibody panel targeting human CD20, CD3, CD163, lysosomal associated membrane protein 3 (LAMP3), programmed cell death protein 1 (PD-1), and von Willebrand factor (vWF) (Supplementary table 3). For nuclei detection, 10⨯ Spectral DAPI (4’6-diamidino-2-phenylindole, SKU FP1490, Akoya Biosciences) was used. 7-plex stainings were acquired at 0.5 µm/pixel on the Vectra Polaris™ Automated Quantitative Pathology Imaging System (Akoya Biosciences) using MOTiF technology. Subsequently, whole slide multispectral image analysis was performed using (i) PhenoChart Whole Slide Viewer (Akoya Biosciences) (ii) InForm® image analysis software (Akoya Biosciences) for spectral unmixing and batch analysis, and (iii) HALO® image analysis software (Indica Labs., United States) for fusing unmixed batched images and downstream analysis. Whole slide quantification was conducted to evaluate the spatial arrangement of CD3+ (T cells) and CD20+ (B cells) populations within the glioma microenvironment. The proximity between CD3+ and CD20+ cells was assessed using a predefined criterion: CD3+ cells located within 50 µm of CD20+ cells. Density heatmaps were generated for CD3+ and CD20+ cells using a radius of 25 µm. Descriptive statistics in R were used to summarize the spatial distribution data of B and T cells.

### Whole exome sequencing and analysis

#### Experimental process

Whole exome sequencing (WES) was performed on 41 samples, inclusive of TLS+ (n=21) and TLS-(n=20) samples. DNA was extracted from FFPE glioblastoma tissue samples using the Maxwell RSC FFPE Plus DNA kit, following the manufacturer’s instructions (Promega). The concentration of DNA was determined using Qubit 3.0 Fluorometer (Thermo Fischer Scientific) and the Qubit dsDNA BR Assay kit (Thermo Fischer Scientific). WES libraries were generated from 750 ng DNA, using the Illumina DNA Prep with Enrichment Kit (Illumina). In brief, genomic DNA was tagmented, cleaned-up and amplified. Clean-ups were performed with AMPureXP beads (Beckman Coulter) at a 1.8 × volume ratio. The cleaned-up libraries were then pooled, hybridized with probes and subsequently amplified. The WES libraries were sequenced on the Illumina NextSeq 1000 platform using NextSeq1000 2 × 100 bp flow cells.

#### Data analysis

Base calls generated by the Illumina Realtime Analysis software were converted into BAM and vcf files using the DRAGEN somatic pipeline on Illumina BaseSpace (Illumina, v3.9.5). The quality control, adapter trimming, and read-level filtering were performed on the DRAGEN Somatic app using the default settings. Sequencing data was aligned to GRCh38/hg38 genome and the tumour mutational burden (TMB) values were calculated within the application. TMB values were imported into R for further statistical analysis. Samples were classified as “TMB-high” when mutation is greater or equal to 10 per megabase^52^ and “hypermutation” when TMB is ≥ 17 mut/Mb^53^.

### Bulk RNA-sequencing and analysis

#### Experimental process

Sixty-four gliomas with and without TLS, matched for IDH-status, age, and gender, were subjected to bulk RNA-sequencing. Total RNA was extracted from 10 ⨯ 10 µm FFPE tissue sections using the Maxwell RSC FFPE RNA kit in accordance with the manufacturer’s manual (Promega, United States). RNA concentration was determined using a Qubit 3.0 Fluorometer using the Qubit RNA BR Assay Kit (Thermo Fischer Scientific, United States). The RNA quality was assessed by the TapeStation system 4150 with the High Sensitivity RNA ScreenTape (Agilent Technologies, United States). RNA libraries were generated from 200 ng RNA using the Illumina total RNA Prep Ligation with Ribo-Zero Plus kit (Illumina, United States). In brief, RNA was depleted of rRNA, followed by fragmentation, cDNA generation, library clean-up and amplification. Clean-ups were performed with AMPureXP beads (Beckman Coulter, United States) with a 1.8 × volume ratio. The cleaned-up libraries were then pooled, hybridized with probes, and amplified. The RNA libraries were sequenced in 100 bp paired-end reads on the Illumina NextSeq 1000 sequencer with NextSeq1000 P2 200 cycles flow cells.

#### Data analysis

Alignment, normalization, and quantification of reads were performed using DRAGEN RNA pipeline (v 4.0.4, Illumina). Human genome version 38 (alt-masked v3, graph enabled) was used as alignment reference. The normalized count matrix was combined with metadata, such as clinical data and run information, in a ‘SummariedExperiment’ object. This object was imported into R for downstream analysis. Long non-coding RNA and pseudogenes were filtered and removed. A total of 21,651 genes were retained for further analysis. Differential gene expression was computed using the R package DESeq2^54^. The differential gene expression model used for DESeq2 included several variables: TLS status, IDH status, sex, batch. Genes with adjusted p (padj) values less than 0.05 and log-fold change of 2 were regarded as significantly differentially expressed. Pathway analysis (GSEA), using the top 20 markers identified in our RNA-seq data, was performed using ReactomePA R package (v 1.9.4)^55^. Pathways with padj values of less than 0.05 were considered to be statistically significant. Cellular deconvolution was performed using SpatialDecon R package^56^ (v 1.12.3) with single-cell data derived from glioma. A custom signature matrix was created to decipher the types of cells present, and this was performed by applying a background noise level of 0.1 to estimate cellular proportions within the two groups (tumors with and without TLS). The matrix was created using a single-cell dataset from glioma, known as CoreGB map^31^, which contains 338,564 cells from 16 datasets encompassing over 100 glioblastoma patients. This reference was also used to highlight cell-type specific gene expression. While the CoreGB map provides detailed annotations for all cell types, we focused on finer annotation levels for specific cell types that are prominent within TLS, including B cells, T cells, NK cells, mural cells, endothelial cells, plasma cells, and fibroblasts. For other cell types, including those within the ‘glial-neuronal lineage’, ‘differentiated-like lineage’, ‘stem-like lineage’, and ‘myeloid lineage’, broader annotation groups were retained. This approach was used to minimize gene spill over and enhance the identification of marker genes for the prominent cell types within TLS. The statistical tests were performed using the Wilcoxon test between the different cell type proportions within each sample using the rstatix (R package v0.7.2). R software (version 4.3) was used to perform all data analysis.

### Spatial transcriptomics using GeoMx^®^ Digital Spatial Profiler (DSP)

#### Experimental process

FFPE tissue cuts (5 µm), taken consecutively from the sames blocks that underwent high-plex immunofluorescence, were placed onto microscope slides and the GeoMx^®^ Human Whole Transcriptome Atlas Assay (NanoString Technologies, United States) was conducted following the manufacturer’s protocol. In brief, slides were baked for 2 hrs for paraffin removal, followed by deparaffinisation and rehydration of tissue sections. Antigen retrieval (Tris-EDTA buffer for 15 mins at 100 C) and proteinase K treatment (1 mg/mL for 15 mins at 37 C) were subsequently performed. Tissues sections were then hybridized with the human Whole Transcriptome Atlas (WTA) probes overnight at 37 C. After two stringent washes (4 ⨯ SSC buffer with formamide at 1:1 ratio) at 37 C for 5 mins, the slides were blocked and incubated with morphology marker antibodies: CD20 (channel 666, 50-0202-82, Invitrogen), CD3 (channel 568, NBP2-54405AF532, Novus Biologicals), CD138 (channel 594, 352324, BioLegend). For nuclei detection, SYTO82^TM^ Orange Fluorescent Nucleic Acid Stain (S11363, Thermo Fisher Scientific) was used. The oligo-barcodes are attached to the fluorescent RNA *in situ* hybridization probes via a photocleavable linker. Tissue sections were loaded into the GeoMx^®^ DSP platform, where the region of interest (ROI) could be selected using the morphology markers. CD20 was used to identify the lymphoid aggregates and regions outside the TLS (tumor area) were selected as control. Due to the size limitation in ROI selection (maximum 660 × 785 µm), multiple ROIs were selected for TLS with bigger region size. Upon the ROI selection, UV light was shined onto the sample by the GeoMx^®^ DSP, releasing the oligo-barcodes for collection and sequencing preparation. Oligonucleotide tags were added with Illumina i5 and i7 dual indexing primers to uniquely index each sample and cleaned up using AMPure XP beads (Beckman Coulter). For library concentration measurement and quality assessment, a Qubit fluorometer (Thermo Fisher Scientific) and a TapeStation System (Agilent Technologies, United States) were used respectively. Sequencing was performed on an Illumina NextSeq 1000 and the GeoMx^®^ NGS Pipeline (v2.0.21) on the Illumina BaseSpace Sequence Hub was used for demultiplexing. A total of 137,308,894 transcripts was generated across 64,638 nuclei, corresponding to an average of 2124.27 transcripts per nucleus.

#### Data analysis

Raw counts from four separate experimental runs were grouped into a single dataset. All further analysis was performed with R version 4.3. The initial processing of raw gene expression count files was conducted utilizing R packages including ‘NanoStringNCTools’, ‘GeomxTools’, and ‘GeoMxWorkflows’. The quality control parameters used in the analysis were as follows: a minimum of 1,000 reads per segment, 80% trimmed reads, 80% stitched reads, 75% aligned reads, 50% sequencing saturation, a minimum of 1 negative control count, a maximum of 9,000 counts in the NTC well, a minimum of 20 estimated nuclei, and a minimum area of 1,000 units. A default value of 2.0 was used as threshold for limit of quantification (LOQ) to eliminate gene segments exhibiting poor signal i.e, if a gene has twice the expression level of the negative probes, it is detected with high confidence. Subsequently, integration of data from biological runs with clinical patient data was performed to generate a SummarisedExperiment object. Normalization of the expression profiles across different runs was accomplished through quantile normalization, followed by the implementation of the ‘Limma’ package to mitigate batch effects. Principal component analysis (PCA) and cluster analysis were subsequently conducted on the spatial data. Clustering was performed using k-means method (k=3) implemented in the ClusterR package^57,58^. Marker genes for each cluster were determined using the *scoreMarker* function of the ‘scater’ package^59^. Similar to the bulk RNA-sequencing data, ‘SpatialDecon’ was employed for cell type deconvolution using the previously mentioned signature matrix. However, the combined use of background value (at 0.01) with nuclei counts obtained from GeoMx DSP, were used to determine the cellular compositions within the three TLS clusters. The background noise values were used in accordance with the recommendations in the package manual.

### Spatial proteome profiling (highplex immunofluorescence) and digital image analysis

The COMET™ spatial proteome profiler (Lunaphore Technologies SA) was employed for highplex immunofluorescence staining and imaging for the detection of 32 markers within a single tissue sample. Nine TLS+ Glioma FFPE sections (3 µm) were generated using the Leica SM 2000R microtome (Leica Microsystems), and incubated at 37 C overnight. Prior to staining, FFPE sections were processed for deparaffinization, rehydration and epitope retrieval using Epredia Dewax and HIER buffer at pH 9 (Lunaphore Technologies SA) and PT Module™ (Epredia), for 60 mins at 102 C. 32-plex sequential immunofluorescence (IF) stainings and digital imaging were performed on the COMET^TM^ platform using an antibody panel for the detection of the following markers: CD3, CD4, CD8, CD20, CD23, CD38, CD68, CD74, CD163, CXCR5, CXCL13, CCL19, αSMA, vWF, ICOS, IL-10, IL-35, IgA, IgG, Ki-67, MZB-1, LAMP3, PD-1, PD-L1, FoxP3, granzyme B (GrzB), GAP43, Col4, Col6A1, decorin (DCN), vimentin (VIM), neurofilament (NF) (Supplementary Table 3). A mixture of Alexa Fluor Plus 555-(1:100, #32732 or #32727, ThermoFisher Scientific) and 647-conjugated (1:200, #32733 or #32728, ThermoFisher Scientific) anti-mouse and anti-rabbit secondary antibodies was applied with DAPI nuclear stain (4’6-diamidino-2-phenylindole, 1:500, #62248, ThermoFisher Scientific). All reagents were prepared in multistaining buffer (BU06, Lunaphore Technologies SA). Multiple, sequential cycles of double-IF stainings and single-channel imaging were followed by the chemical elution of primary and secondary antibodies using an elution buffer (BU07-L, Lunaphore Technologies). Autofluorescence was reduced both, chemically by applying the Quenching buffer (BU08-L, Lunaphore Technologies) and digitally, by software-based background subtraction (HORIZON™, Lunaphore Technologies). TIFF output images was performed using the HALO^®^ image analysis platform (Indica Lab). Infiltration analysis of CD20+ B cells, CD4+ and CD8+ T cells and IgG+MZB1+CD38+ or IgA+CD38+ plasma cells was performed around the Col6A1 interface at the distance of 200 µm inside TLS and 200 µm outside TLS and Col6A1 network.

R Statistical Software v4.3 was used for statistical analysis and displaying results. For this, the results from cell segmentation (x-y coordinates of each cell object and marker expression) were exported from HALO. Cellular phenotypes were determined according to Supplementary Table 2. For each cell, we calculated the mean distance to the closest three B cells. Cells for which this distance was <600 µm were considered inside TLS, since this allows us to segregate between cells that are located within and close to B cell aggregates.

