## Supplementary_Table_3 for "Glioma-associated tertiary lymphoid structures are sites of lymphocyte clonal expansion and plasma cell formation"

List of primary antibodies applied in 32plex sequential-IF and 7plex Opal-TSA panels

| Marker | Source | Dilution  32plex Seq-IF (COMET™) | Dilution  7plex (LabSat®) | Dilution  Opal-TSA |
| --- | --- | --- | --- | --- |
| αSMA | DAKO Agilent, M0851 | 1:500 |  |  |
| CCL19 | Thermo Fisher Scientific, MA5-26657 | 1:800 |  |  |
| CD3 | DAKO Agilent, A0452 | 1:150 | 1:450 | Opal-780 1:25 |
| CD4 | Thermo Fisher Scientific, MA5-16338 | 1:50 |  |  |
| CD8 | DAKO Agilent, M7103 | 1:100 |  |  |
| CD20 | DAKO Agilent, M0755 | 1:350 | 1:120 | Opal-570 1:120 |
| CD23 | Abcam, ab16702 | 1:100 |  |  |
| CD38 | GeneTex, GTX01959 | 1:50 |  |  |
| CD68 | DAKO Agilent, M0876 | 1:50 |  |  |
| CD74 | Abcam, ab9514 | 1:120 |  |  |
| CD163 | Abcam, ab265592 | 1:700 | 1:500 | Opal-620 1:120 |
| Col4 | Abcam, ab214417 | 1:200 |  |  |
| Col6A1 | Abcam, ab151422 | 1:250 |  |  |
| CXCL13 | Abcam, ab246518 | 1:400 |  |  |
| CXCR5 | R&D Systems, MAB190R | 1:150 |  |  |
| DCN | Abcam, ab268048 | 1:100 |  |  |
| FoxP3 | Thermo Fisher Scientific, MA5-16365 | 1:50 |  |  |
| Gap43 | Abcam, ab75810 | 1:2000 |  |  |
| GrzB | Abcam, ab298586 | 1:150 |  |  |
| ICOS | Abcam, ab224644 | 1:100 |  |  |
| IgA | Abcam, ab124716 | 1:60 |  |  |
| IgG | Abcam ab109489 | 1:400 |  |  |
| IL-10 | GeneTex, GTX632359 | 1:50 |  |  |
| IL-35 | My Biosource, MBS2003070 | 1:100 |  |  |
| Ki-67 | DAKO Agilent, M7240 | 1:40 |  |  |
| LAMP3 | Abcam, ab271053 | 1:500 | 1:500 | Opal-690 1:150 |
| MZB1 | Thermo Fisher Scientific, MA5-29430 | 1:800 |  |  |
| NF | DAKO Agilent, M0762 | 1:500 |  |  |
| PD-1 | Abcam, ab137132 | 1:250 | 1:300 | Opal-520 1:100 |
| PD-L1 | Abcam, ab228415 | 1:500 |  |  |
| VIM | DAKO Agilent, M0725 | 1:500 |  |  |
| vWF | DAKO Agilent, A0082 | 1:120 | 1:120 | 1:150 Opal-480 |
